# Programmatic goals and spatial epidemiology influence the merit of targeted versus of population-wide interventions for yaws eradication

**DOI:** 10.1101/640326

**Authors:** Eric Q. Mooring, Michael Marks, Oriol Mitjà, Marcia Castro, Marc Lipsitch, Megan B. Murray

## Abstract

Infectious disease eradication programs often pursue spatially targeted interventions, but how well they perform might depend on the underlying spatial epidemiology and the specific goal of the program. We use a stochastic compartmental metapopulation model of yaws transmission to investigate how total targeted treatment (TTT) performs compared to mass drug administration (MDA) in different settings. While TTT can efficiently control the prevalence of active yaws disease, we consistently found that multiple rounds of TTT are required to match the impact of 1 round of MDA on the prevalence of latent yaws infection. When complete eradication of yaws is the goal, MDA can achieve the same result as TTT more quickly and probably at lower cost. We found that the performance of TTT is improved when there is little mixing between subpopulations and when there is spatial heterogeneity in transmissibility, but even in these settings, our model suggests that MDA will still outperform TTT.

**Significance Statement:** Yaws is a neglected tropical disease that causes skin lesions. Eradicating yaws is challenging because people can be infected but not show any signs or symptoms for years. Using simulations, we found that targeting antibiotics to people with active yaws and to their neighbors is a good way to combat active yaws, but treating entire populations is a better way to get rid of all infections, including the hidden ones. Also, targeted treatment works better in populations in which people do not move around much and in which how easily the disease is transmitted varies from place to place. Overall, a targeted treatment strategy uses fewer antibiotics but takes longer than mass treatment to achieve results.

## Introduction

Public health officials aiming to eradicate a disease often must choose between implementing an intervention throughout an entire population and targeting the intervention to a subgroup. In this study we focus on one example of this choice: how to optimally deploy antibiotic treatment to eradicate yaws. Yaws is a neglected tropical disease (NTD) caused by the bacterium *Treponema pallidum* subspecies *pertenue*. The disease causes skin lesions and arthralgia and can be disfiguring if left untreated. Like other treponemal infections, the infection can persist in an asymptomatic latent stage which can subsequently reactivate and become symptomatic and infectious [1]. Eradicating yaws therefore will involve the treatment of both active yaws and latent infection.

The World Health Organization’s yaws eradication strategy calls for 1 or more rounds of mass drug administration (MDA) with oral azithromycin. During MDA, all people living within a defined area are offered treatment regardless of disease status. MDA is then followed by episodic rounds of total targeted treatment (TTT) in which the entire population is screened for active yaws, and then only patients with suspected active yaws and their contacts are treated [2]. Field demonstration projects as well as transmission models have suggested that a single round of MDA followed by multiple rounds of TTT is insufficient to achieve yaws eradication [3, 4]. The optimal sequence of rounds of MDA and rounds of TTT is an unresolved challenge [5].

The underlying spatial epidemiology may influence how well TTT performs compared to MDA. For example, targeted treatment may perform better when small communities are less interconnected. Also, spatial heterogeneity in transmissibility may influence how well TTT performs relative to MDA; targeting interventions to locations most in need may help avoid excessive treatment in areas in which the disease is less able to persist. (Because yaws risk factors such as poor hygiene, overcrowding, and a moist environment vary spatially, spatial heterogeneity in transmissibility is plausible and may be among the causes of observed small-scale spatial heterogeneity in yaws prevalence [6, 7].) Finally, we study how well TTT can prevent imported yaws cases from re-establishing the disease in an area.

The choice between population-wide versus targeted interventions also depends on the relative costs of the different approaches [8]. In settings that are relatively inaccessible, the multiple rounds of MDA strategy will require more drug but may cost less overall if it reduces the number of times that communities are visited. Conversely, in more accessible populations, drugs may make up a larger proportion of program costs and repeated rounds of TTT may be preferred.

In this study, we assess whether and how TTT and MDA strategies for yaws depend on the spatial epidemiological context as well as on the specific goal of the program. Specifically, we investigate the consequences of population mixing between communities, spatial heterogeneity in transmissibility, and imported yaws cases.

## Results

We simulated yaws transmission (Figure S1) and intervention campaigns in populations averaging 20000 people divided among 200 “hamlets” that ranged in size from 75 to 125 people. We assumed an average MDA coverage of 90%. Typically, TTT has entailed treating just household and school contacts of patients with suspected active yaws, but analysis of serological data has suggested that it might be better to use a broader definition of contact [9]. Therefore, we defined contacts to be anyone living in the same hamlet as a case of active yaws. To model TTT, we assumed that cases of active disease were detected with probability of 90% and that treatment was administered to, on average, 90% of all persons living in any hamlet with at least 1 detected active case (Table S1).

We measure how well TTT performs versus MDA by using simulations to calculate the amount of TTT required to meet or exceed the impact of 1 additional round of MDA, given the underlying spatial epidemiology and given the number of initial rounds of MDA (either 1, 2, or 3 initial rounds) (Figure 1). We consider 2 indicators of the impact of the additional round of MDA: 1) the prevalence of active disease and 2) the prevalence of latent infection. Both indicators are measured 6 months after the additional round of MDA. The amount of TTT is measured in 2 dimensions: 1) the number of rounds of TTT and 2) the total number of sets of contacts treated.

**Figure 1.**
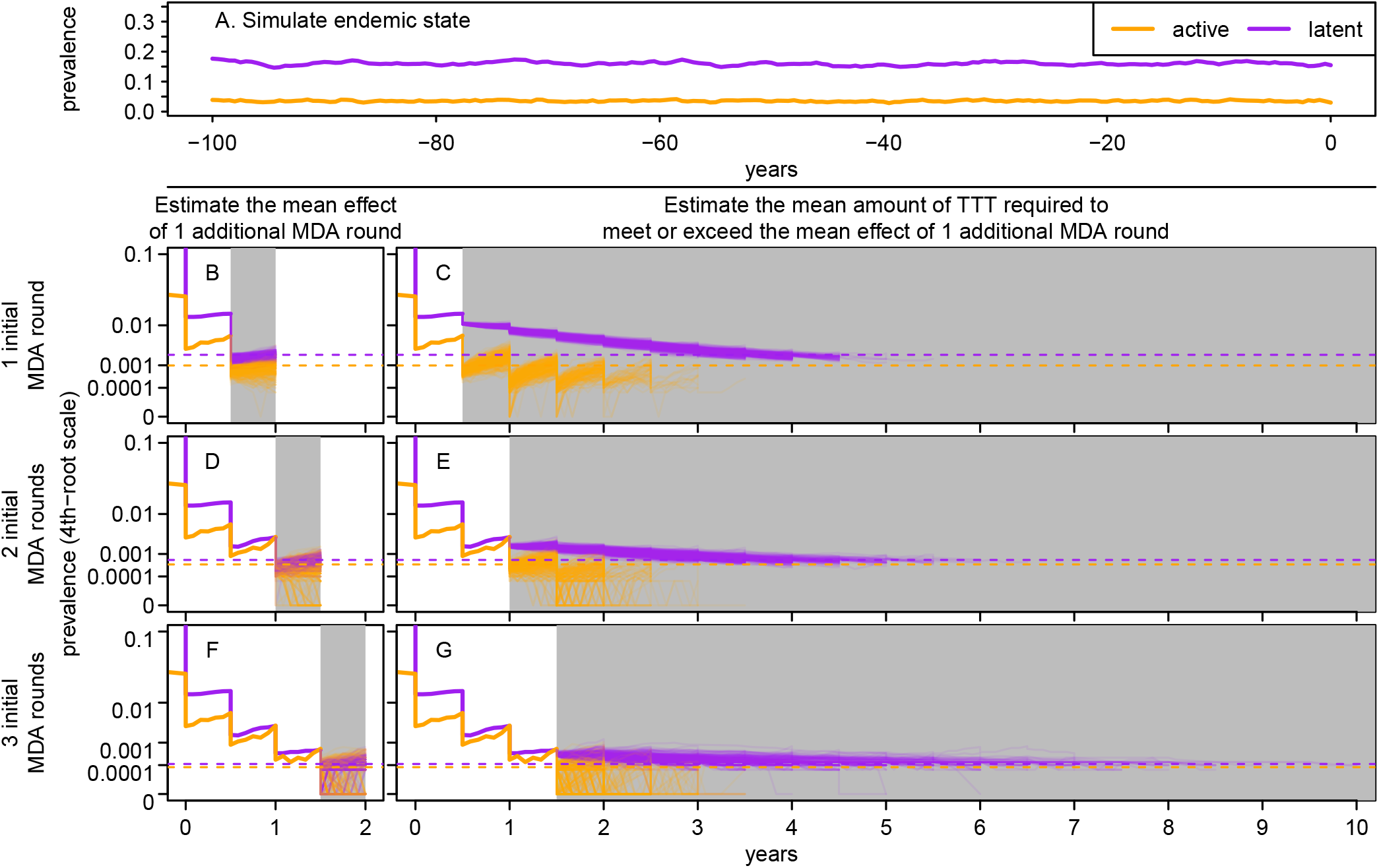
Illustration of TTT versus MDA comparison method. Simulation of an endemic state (A) was followed by up to 3 initial rounds of MDA to develop 3 sets of baseline states. For each baseline state, an additional round of MDA (grey shaded regions of B, D, F) was simulated 300 times to estimate its average impact. Purple and orange dashed lines indicate the mean prevalence of latent infection and active disease, respectively, following the additional round of MDA. For each baseline state we simulated 300 TTT campaigns (grey shaded regions of C, E, G) to estimate the average amount of TTT required to match the average impact of the additional MDA round (grey shaded regions of B, D, F). In this example, mixing between hamlets and spatial heterogeneity in transmissibility are set at intermediate levels.

To model different spatial epidemiological settings, we varied the strength of the epidemiological connections between hamlets: We considered scenarios where 71% (high amount of mixing), 39%, or 14% (low amount of mixing) of contacts occurred between hamlets (Figure 2A). We explored 3 different levels of spatial heterogeneity in transmissibility at the hamlet-scale (Figure 2B) as well as scenarios in which there was no exogenous importation of yaws into the population or where the importation rate corresponded to an average of 10 active cases imported per year into a wholly susceptible population of 20000 people.

**Figure 2.**
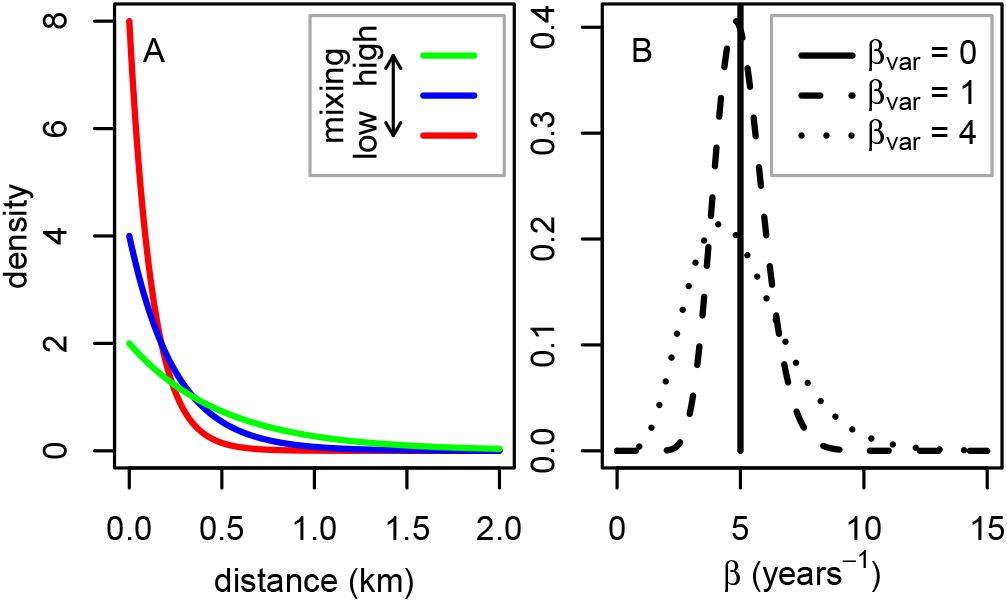
Probability distributions used to model mixing between hamlets (A) and spatial heterogeneity in transmissibility (B).

### Utility of TTT versus MDA depends on indicator of progress toward eradication

When the indicator of progress towards eradication is solely the prevalence of active infection, TTT may be as costly or less costly than MDA (Figure 3A-C). This is especially true when TTT is compared to MDA after 2 or 3 initial rounds of MDA; in most simulations only a few hamlets need to be treated over the course of 2 to 3 rounds of TTT in order to match or exceed the impact of an additional round of MDA on the prevalence of active infection (Figure 3B, C). In other words, if the goal of a program is solely to reduce the prevalence of active disease, then 2 to 3 rounds of MDA followed by 2 to 3 rounds of TTT is generally equivalent to 3 to 4 rounds of MDA. Two to 3 rounds of TTT in which few doses of azithromycin are distributed may cost less than 1 additional round of MDA, especially in geographic settings that are relatively accessible.

However, TTT does not lead to as rapid of reductions in the prevalence of latent infection. While repeated TTT is eventually able to match the impact of a round of MDA on the prevalence of latent infection, our results suggest that in many scenarios, 5 or more rounds are required to match a single round of MDA (Figure 3D-F). Our results almost always fall far beyond the threshold where MDA costs as much as TTT. A single round of MDA is almost certain to cost less than the numerous rounds of TTT that on average are required to match its impact on the prevalence of latent infection. A similar result would be found if the indicator were prevalence of all infections combined, because most infections are latent rather than active.

**Figure 3.**
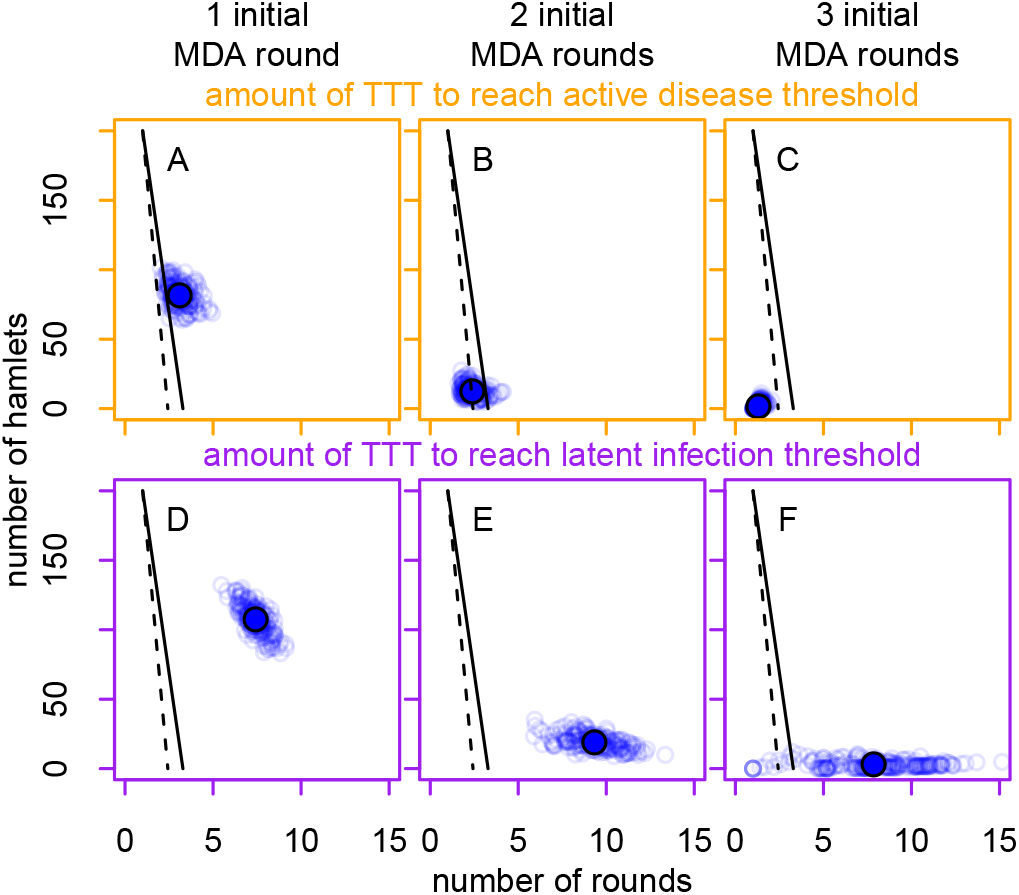
Number of rounds of TTT and total number of hamlets that must be treated across those rounds in order to match the impact of a single round of MDA. The indicator of intervention impact is prevalence of active disease in the row of panels with orange borders (A-C) and is prevalence of latent infection in the row of panels with purple borders (D-F). The columns correspond to the initial number of rounds of MDA. The large solid colored circles mark the mean result, while the small circles correspond to means calculated from different stochastically generated baseline states under fixed parameters. The solid and dashed black lines correspond to the thresholds where TTT and MDA cost the same and represent low- and high-operational-cost scenarios, respectively. TTT is preferable in scenarios to the left/below each line and MDA is preferable in scenarios to the right/above each line. In this figure, mixing between hamlets and spatial heterogeneity in transmissibility are set at intermediate levels.

### Consequences of varying mixing between hamlets and varying spatial heterogeneity in transmissibility

Figure 4 shows how varying different aspects of spatial epidemiology affects the relative performance of TTT versus MDA. Here we focus on prevalence of latent infection as the indicator used to compare TTT to MDA. (Full results for both indicators are given in Figure S4.) The performance of TTT is improved when hamlets are less connected, especially when TTT is compared to MDA after just 1 initial round of MDA. When hamlets are less connected, only about a third to half as many hamlets must be treated over the course of the campaign (Figure 4A, D, G). One reason for this pattern is that when hamlets are less connected, the equilibrium prevalence of infection is lower. (With less mixing, we expect more frequent stochastic fadeouts.) However, this does not explain the entire pattern; when mixing is less, cases are likely to be clustered within fewer hamlets, thereby making TTT more efficient. Notably, when there is no spatial heterogeneity in transmissibility, less mixing between hamlets on average slightly increases the number of rounds of TTT required to match the impact of a round of MDA on the prevalence of latent infection (Figure 4A). When hamlets are more connected, we expect that active cases are more frequently seeded in hamlets that have not been recently treated, which would trigger the treatment of those hamlets and the cure of latent infections, including latent infections that were already present and are not directly epidemiologically linked to the active case that triggered treatment. (This cannot be confirmed empirically in our model, however, because the compartmental model does not explicitly model transmission chains and therefore we cannot distinguish between active disease due to infection from outside the hamlet and active disease due to reactivation of latent infection.)

**Figure 4.**
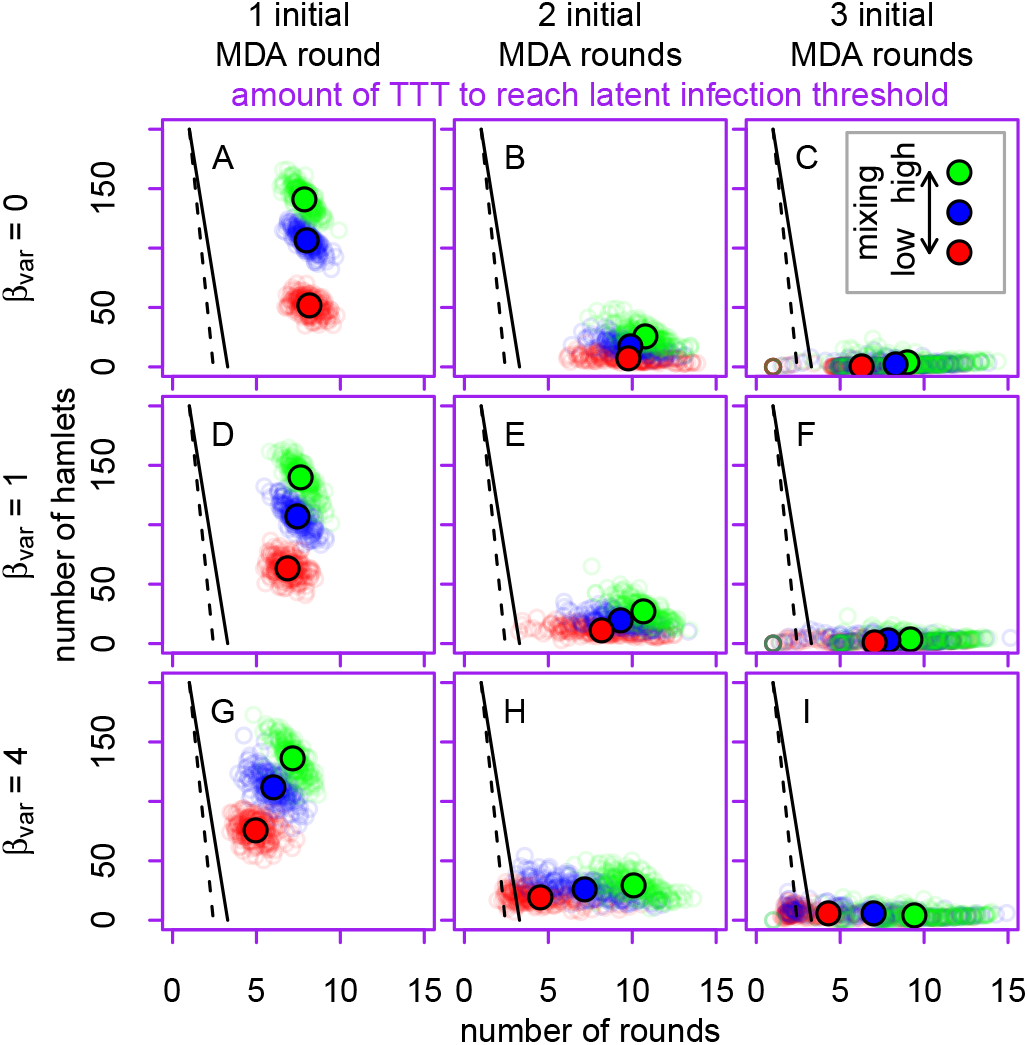
Number of rounds of TTT and total number of hamlets that must be treated across those rounds in order to match the impact of a single round of MDA. The colors of the points correspond to mixing between hamlets, the rows to spatial heterogeneity in transmissibility, and the columns to the initial number of rounds of MDA. The large solid colored circles mark the mean result for each set of parameters, while the small circles correspond to means calculated from different baseline states. The solid and dashed black lines correspond to the thresholds where TTT and MDA cost the same and represent low- and high-operational-cost scenarios, respectively.

When TTT is compared to an additional round of MDA following 3 initial rounds of MDA we again observe that TTT’s performance improves when mixing between hamlets is less (Figure 4C, F, I). But the specifics of our results differ: Regardless of mixing, after 3 initial rounds of MDA the total number of hamlets requiring treatment during TTT is very low, but when mixing is reduced, the number of rounds of TTT is lower. The pattern following 2 initial rounds of MDA is intermediate between the pattern after just 1 round and the pattern after 3 rounds.

Increasing spatial heterogeneity in transmissibility reduces the number of rounds of TTT required to match the impact of an additional round of MDA (Figure 4), but the reduction is not large enough to make TTT clearly superior to MDA. The effect of spatial heterogeneity in transmissibility was most pronounced when mixing between hamlets was low. The greater the mixing between hamlets, the more transmission dynamics merely reflect average transmissibility across the entire population. Sources of variability in our results are discussed in the Supporting Information (Figures S5–S8).

### Performance of TTT when yaws is imported from outside the study area

The introduction of cases of yaws from outside the modelled population does not dampen the ability of TTT relative to MDA to reduce the prevalence of active disease (Figure S9 A-I). After a single initial round of MDA, introduced cases do not seem to substantially affect how TTT performs versus MDA at reducing the prevalence of latent infection (Figure 5A, D, E). But introduced cases become more consequential as the overall prevalence of yaws declines. After 2 or 3 initial rounds of MDA, a TTT campaign may not be able to match the impact of an additional round of MDA on the prevalence of latent infection (Figure 5B, C, E, F, H, I). Especially when mixing between hamlets is high, the probability that a TTT campaign has matched the impact of a single additional round of MDA on the prevalence of latent infection may remain less than 50% even after 30 rounds of TTT. If introduced cases of active yaws become latent before they can be identified during biannual rounds of TTT, they can persist as latent cases if there are not active cases to otherwise trigger treatment in these hamlets.

**Figure 5.**
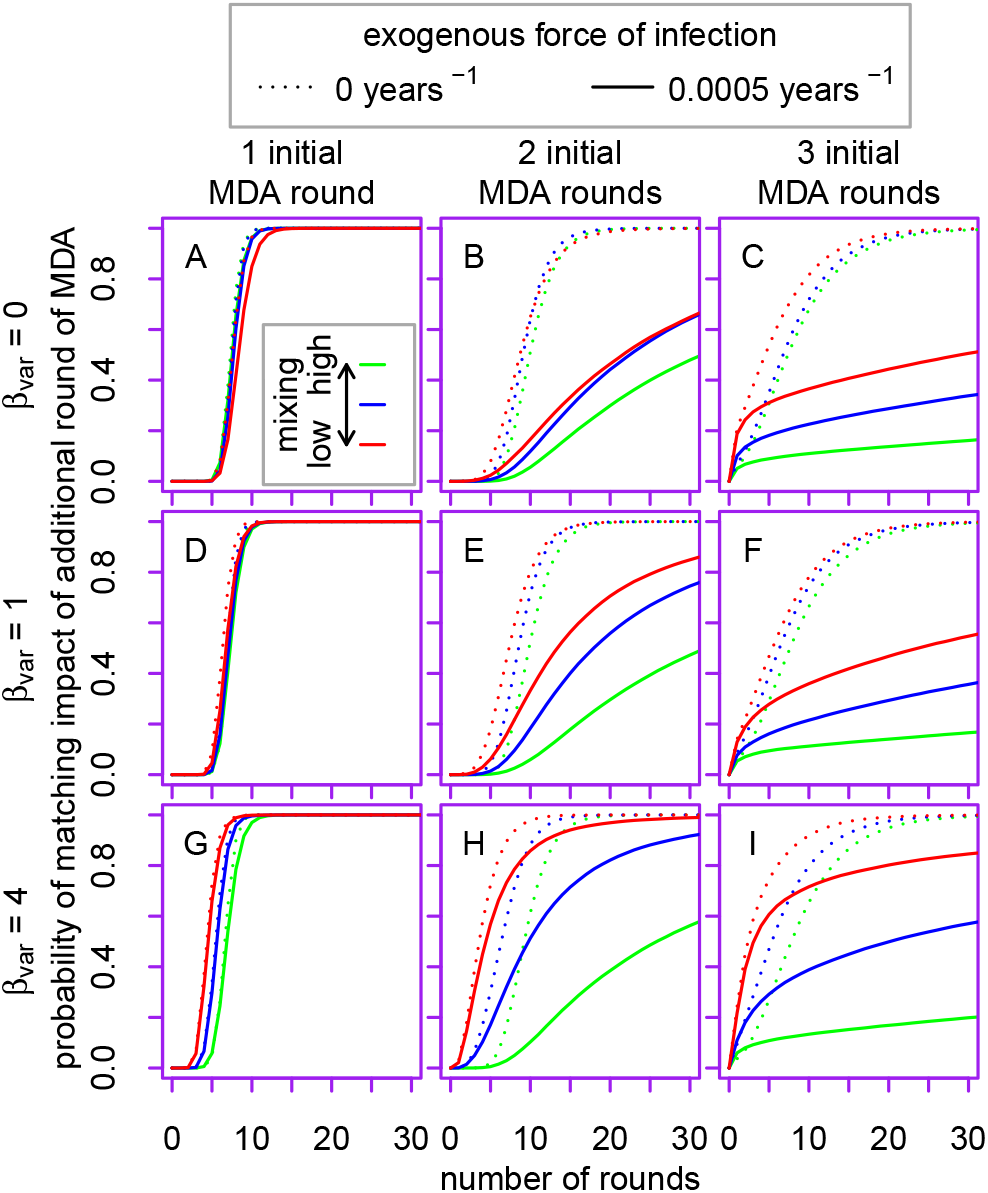
Effect of an exogenous force of infection on the probability of TTT to match the impact of an additional round of MDA. The panels plot the probability of matching the impact of an additional round of MDA as a function of number of rounds of TTT, when the indicator of impact is prevalence of latent infection. The colors of the lines correspond to the mixing between hamlets, the rows to spatial heterogeneity in transmissibility, and the columns to the initial number of rounds of MDA.

### Changing the frequency of TTT

When TTT is quarterly rather than biannual, the number of rounds of TTT required to match the impact of MDA on the prevalence of latent infection approximately doubles (Figures 6, S10). This suggests that declines in latent infection are primarily tied to the passage of calendar time rather than to the number of rounds of treatment. TTT prevents yaws from rebounding while the number of latent cases at risk of reactivating declines with the passage of time.

## Discussion

Across numerous spatial epidemiology scenarios, our models consistently show that MDA is a powerful tool to reduce the prevalence of latent yaws infection. While choosing TTT over MDA reduces demand for azithromycin, the eradication campaign requires repeated rounds of TTT. Even in settings relatively favorable to TTT, the costs involved in repeatedly screening the population likely outweigh the savings accrued from using less azithromycin. This finding aligns with the results from a previous transmission model and with a field trial from Papua New Guinea where an initial round of MDA greatly reduced the prevalence of active yaws but occasional cases of yaws still occurred even after numerous subsequent rounds of TTT [3, 4].

The performance of TTT relative to MDA depends on the underlying spatial epidemiology. The specific consequences of changing the mixing between hamlets depends on the number of initial rounds of MDA, but all else equal, the performance of TTT is improved when hamlets are less connected. Similarly, TTT’s performance improves if there is spatial heterogeneity in transmissibility, though the specifics of the effect depend on the amount of mixing between hamlets. Sustained TTT is poorly suited to reducing the prevalence of latent infection in the face of low-level but sustained importation of yaws from outside the population.

**Figure 6.**
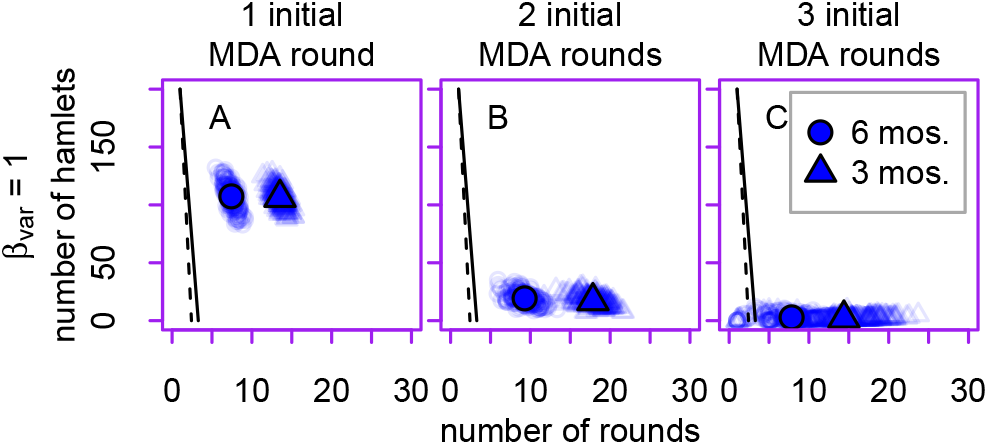
Comparison of quarterly (triangles) versus biannual (circles) rounds of MDA and TTT on the amount of TTT required to match the average impact of an additional round of MDA on the prevalence of latent infection. Intermediate levels of spatial heterogeneity in transmission and in mixing are illustrated here.

Our model is not designed to make quantitative predictions for any real-world location. But insights from this study can help inform yaws eradication strategy: This study supports the view that MDA—and especially repeated rounds of MDA—is a tool for reaching eradication more quickly and likely at lower cost, rather than a tool that is absolutely required to reach eradication. Indeed, in the pre-azithromycin era, mass treatment was not always a part of yaws eradication strategies, and our findings are compatible with observational data showing that sustained targeted treatment programs can achieve results [10]. TTT is potentially more feasible in settings in which transmission is more spatially constrained, but even when we simultaneously modelled low levels of mixing and high spatial variability of transmissibility, multiple rounds of TTT were typically required to match the impact of a single round of MDA. Therefore, in most settings our model predicts that MDA is likely to be a less expensive way to reduce the prevalence of yaws.

We have focused on the average performance of TTT versus MDA in various circumstances. We have not focused on modeling campaigns all the way to eradication and therefore we do not directly address when to stop implementing both MDA and TTT. In some simulations of the model, 3 initial rounds of MDA achieved eradication. In those instances, an additional round of MDA would be superfluous. The central challenge for public health officials is to know if eradication has, in fact, been achieved. It is important to note that mass screening is an essential part of TTT and provides critical information on progress to eradication. In fact, when close to achieving eradication, rounds of TTT can be thought of as rounds of mass screening to confirm eradication in which public health workers are also prepared to treat any cases (plus contacts) that might still exist in the population.

It is overly simplistic to see yaws eradication as a linear path from MDA to TTT to surveillance to eradication. Detecting occasional cases of yaws across multiple rounds of TTT may signify that there are still latent infections in the population and that a round of MDA might be warranted. Alternatively, public health officials may decide to just keep doing TTT recognizing that while this choice requires few doses of azithromycin, it probably will delay achieving eradication.

Our study is limited by uncertainty in model structure and parameters. Importantly, latent infection is not well understood. Not only is the average duration of latent infection unclear, but the distribution of these durations is unknown. This model implicitly assumes that the duration of latent infection follows an exponential distribution: a small fraction of latently infected people remains at risk of reactivating for a long period of time. If in the real world the period during which latent infections are at risk of reactivation is of shorter duration than modelled here, then this study probably understates the utility of TTT. Additionally, we assume that there is no systematic non-adherence. If some people are consistently missed by a treatment campaign, then the impact of MDA can be substantially lessened [11]. But TTT may well suffer from a similar limitation. Further research should study systematic non-adherence to targeted interventions.

We did not model a continually operating surveillance system that treats contacts whenever a case of active yaws is found, nor did we consider whether a high-quality surveillance system can counteract low-coverage MDA or TTT and vice versa. Attaining a coverage level of 90% is difficult; programs have generally not reached this coverage level even in field demonstration projects [12]. Future modelling work should explore overall program performance considering varying amounts and intensity of mass treatment, targeted treatment, and continual surveillance and treatment.

Our economic model was quite simple. We did not account for drug resistance [13] nor any non-yaws-related health benefits that may accrue from MDA [14]. Also, we assumed that costs were a function of the number of rounds, not calendar time. But some costs (e.g. maintaining community-based surveillance and program administration) are relatively constant per year, regardless of the number of rounds of MDA or TTT implemented. Consequently, less frequent TTT may be less attractive than our results suggest. Finally, we do not consider synergies that may arise from integrated skin NTD control programs [15]: If health systems were to commit to ongoing high-quality dermatologic screening in order to combat diseases other than yaws, then a TTT-based yaws eradication strategy might be preferable. On the other hand, integration of yaws programs with other NTD programs that implement recurrent cycles of preventive chemotherapy would favor the MDA approach.

Understanding the relative merits of targeted versus population-wide infectious diseases control interventions is complex and ill-suited to generalizations. We have shown that how well targeted interventions perform depends on multiple and interacting components of spatial epidemiology. Disease natural history is another key consideration when comparing targeted versus population-wide interventions: In the case of yaws, MDA is much better than TTT at quickly removing latent infections. But for diseases without a long latent stage, targeted interventions may be ideal: Targeted interventions such as vaccination can be highly effective for acute epidemic-prone infectious diseases [16, 17, 18]. A key part of the cost of TTT is the cost of repeatedly screening the population even when there are hardly any active cases. Consequently, targeted interventions are comparatively better suited to disease that are likely to come to the attention of health providers even in the absence of active case detection (e.g. because of clinical severity or unique signs and symptoms).

Interest in targeted interventions for a wide range of infectious diseases—including malaria and tuberculosis—is likely to only grow [19, 20]. Each disease and each geographic setting presents its own challenges and opportunities for targeted interventions. Researchers and public health practitioners considering spatially targeted infectious disease interventions must be explicit about their goal. A targeted strategy may be an efficient way to control the burden of disease or to achieve elimination as a public health problem but may not be as well-suited for an eradication program.

## Methods

Our transmission model is based on a previously published yaws model that we modified to include births and deaths and an exogenous force of infection [4]. We also modified it to be a metapopulation model [21]. Hamlets were randomly located within a square area of 100 km^2^ and we assume that the strength of coupling declines exponentially with increasing distance. The model was implemented using the adaptive tau-leaping approximation to the Gillespie algorithm [22, 23, 24]. Initial conditions for the stochastic model were set at equilibria values from a deterministic version of the model [25]. Full details of the model are given in the Supporting Information. For each number of initial rounds of MDA (1, 2, or 3) within each baseline simulation, we estimated the average impact of an additional round of MDA by replicating an additional round 300 times. We then simulated 300 TTT campaigns to estimate the amount of TTT required to match the average impact of that additional round. TTT campaigns consisted of a minimum of 1 and a maximum of 100 rounds of TTT (Table S5). We conducted 200 independent simulations at each of 18 combinations of spatial epidemiology parameters (3 levels of coupling, 3 of spatial heterogeneity of transmissibility, and 2 of exogenous force of infection) with biannual MDA and TTT. We also modelled quarterly MDA and TTT. To create our cost scenarios, we consulted the published literature on yaws interventions and other mass drug administration campaigns [26, 27]. The cost of implementing MDA is well documented, but TTT-related costs are more speculative (Table S6). We calculated for each scenario the threshold where the cost of a TTT campaign matches the cost of a round of MDA.

## Acknowledgements

EQM was supported by T32 AI 007535 from the National Institute of Allergy and Infectious Diseases. The computations in this analysis were run on the Odyssey cluster supported by the FAS Division of Science, Research Computing Group at Harvard University.

## Supporting information

### Modelling yaws natural history

We modified a previously published compartmental model structure of yaws [1] to include births and deaths at rate *b* and an exogenous force of infection at rate *ξ* (Figure S1). In the model, people are born uninfected and susceptible to yaws (compartment *S*) and become infected after exposure to an infectious patient. Once infected, they enter the primary active yaws stage which is both symptomatic and infectious (compartment *I*_1_) from which they can either 1) be cured and return to the susceptible stage, 2) die, 3) move into a secondary active stage (also symptomatic and infectious) (compartment *I*_2_), or 4) move into an asymptomatic latent infection stage which is not infectious (compartment *L*). They then move back and forth between these 2 stages until they or are cured, at which point they reenter the susceptible stage, or die. The model assumes that latently infected individuals can develop secondary active yaws but cannot be re-infected.

**Figure S1:**
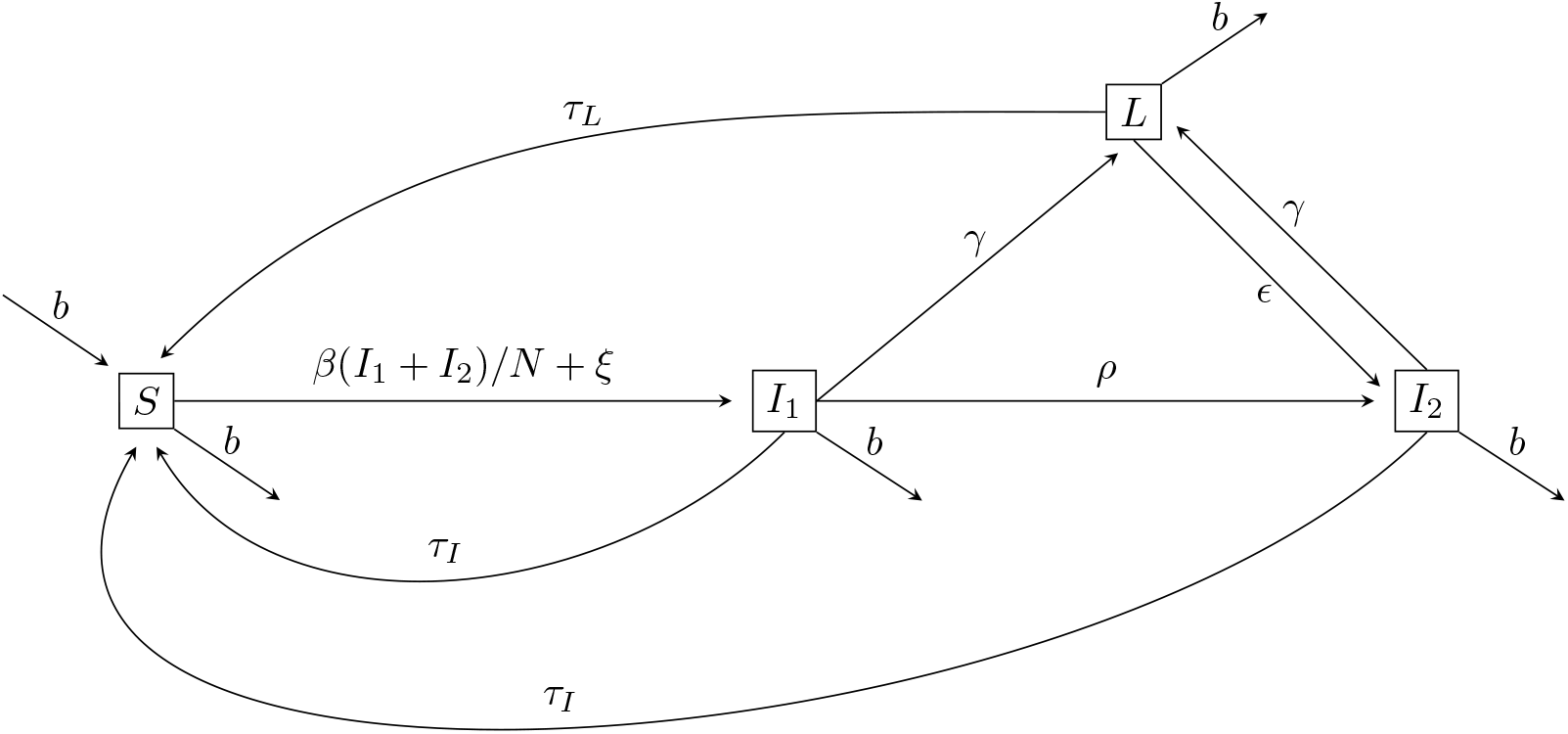
Box-and-arrow diagram of compartmental model that underpins all simulations. The parameters are defined in Table S1. Compartments are susceptible (*S*), primary active yaws (*I*_1_), secondary active yaws (*I*_2_), and latent infection (*L*).

**Table S1:**
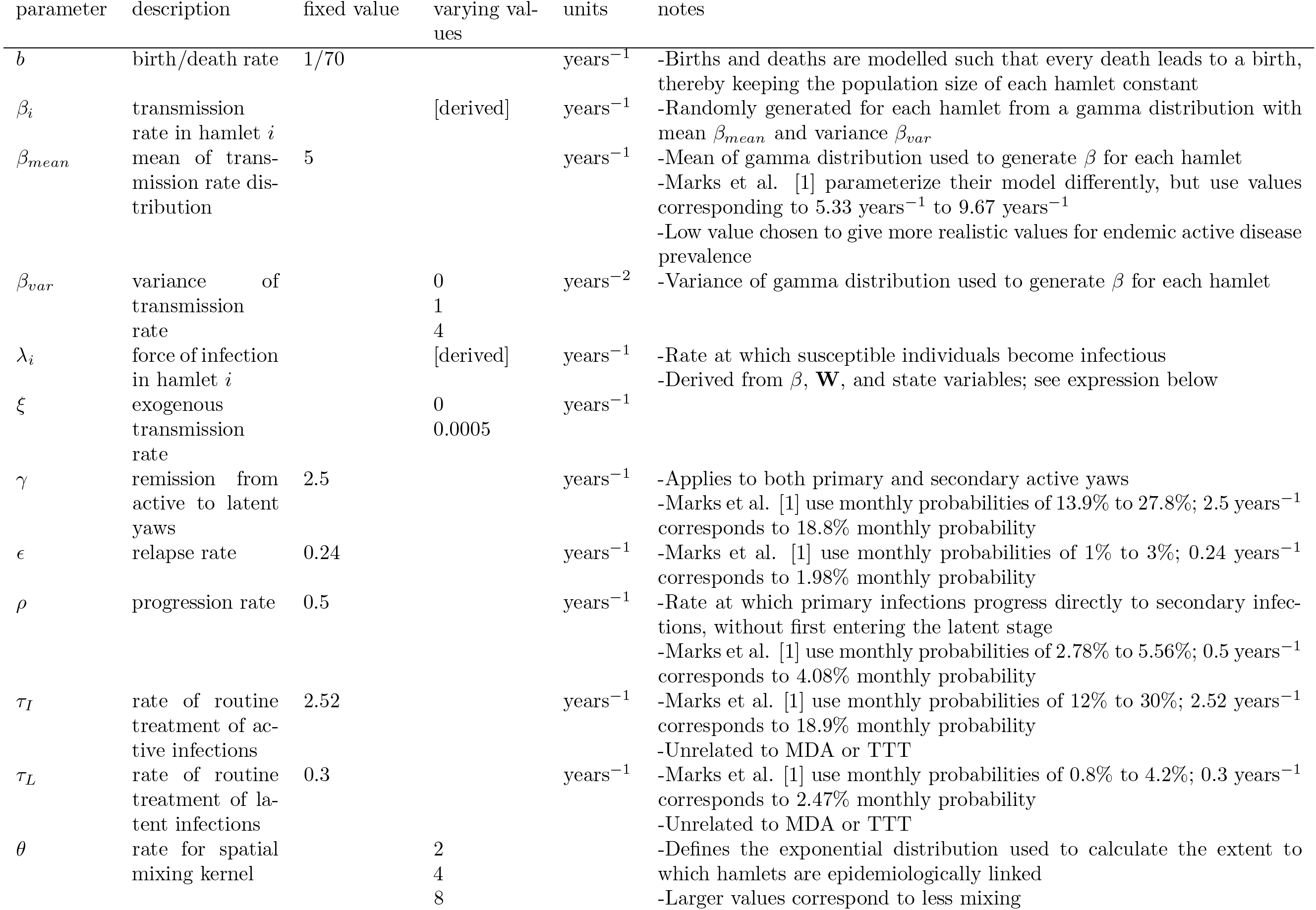
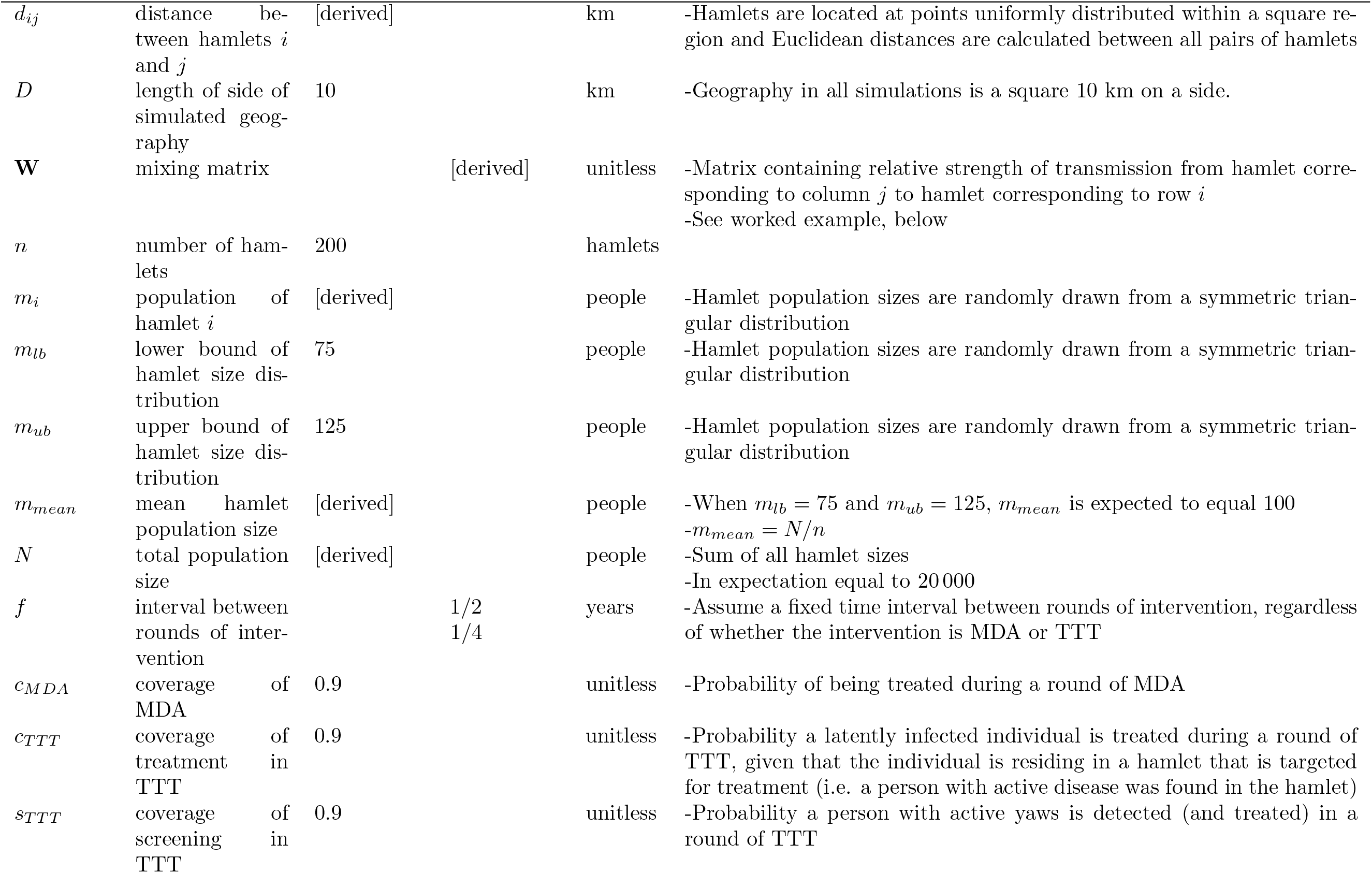
Definitions and values of parameters. (Additional parameters describing costs are given in Table S6.)

### Modelling spatial transmission

We created a metapopulation model with 200 subpopulations which we refer to as hamlets. The population of each hamlet is randomly drawn from a symmetric triangular distribution with a minimum population of 75 and a maximum population of 125. Therefore, the entire population size is, in expectation, 20 000 people. The hamlets are randomly distributed within a square of area 100 km^2^. Spatial transmission is modelled through the following expression:

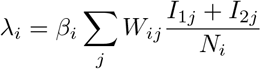

Where *β*_*i*_ is the transmission parameter in hamlet *i*, and *W*_*ij*_ is an element in matrix **W**, which defines mixing; specifically, it is the relative strength of transmission to hamlet *i* from hamlet *j*. In this formulation, yaws spreads across hamlets when infectious individuals move from their hamlet, infect others, and then return home [2]. We assume that the strength of mixing between hamlets declines exponentially with increasing Euclidean distance. The following example with 10 hamlets in a square area of 5 km^2^ illustrates how the mixing matrix **W** is calculated (Figure S2). The following expression defines the exponential kernel used to obtain relative weights between hamlets:

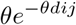

In this example, we assume that *θ* = 4 (Figure S3).

**Figure S2:**
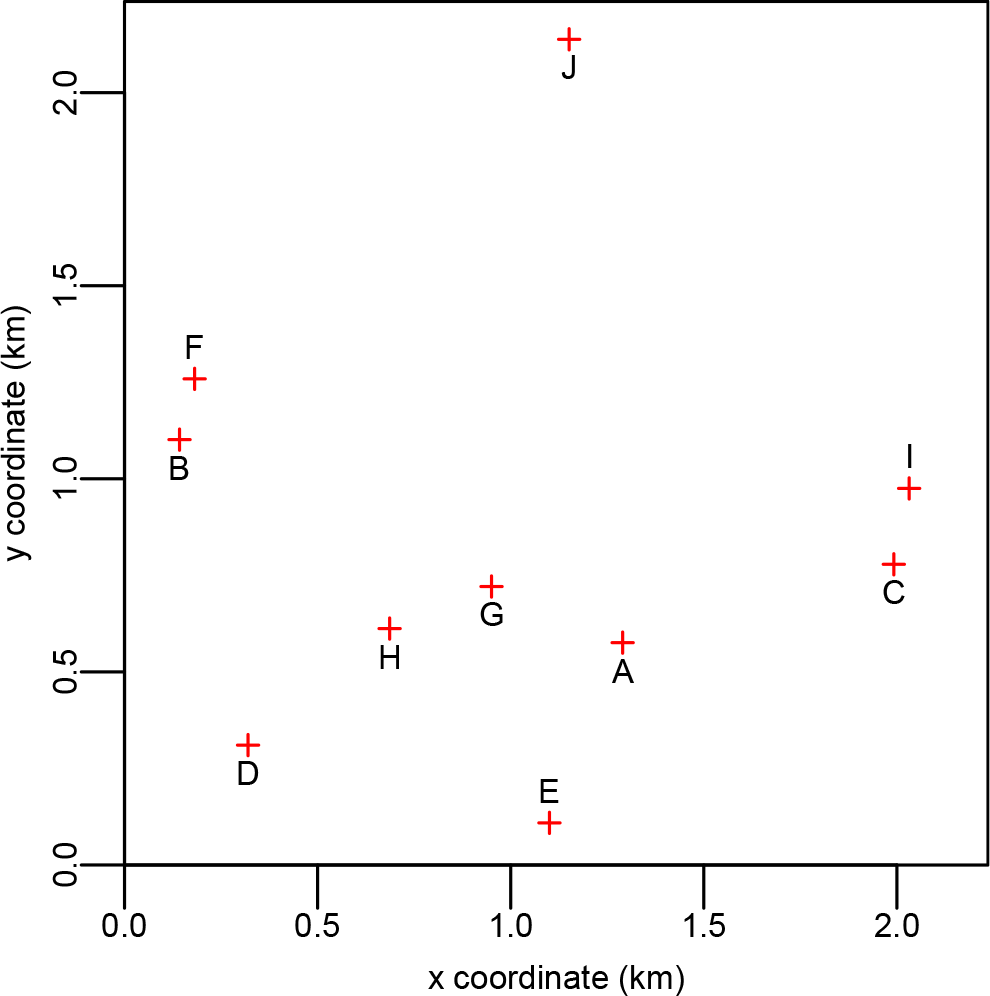
Plot of 10 simulated hamlet locations (A-J) in a 5 km^2^ area.

**Figure S3:**
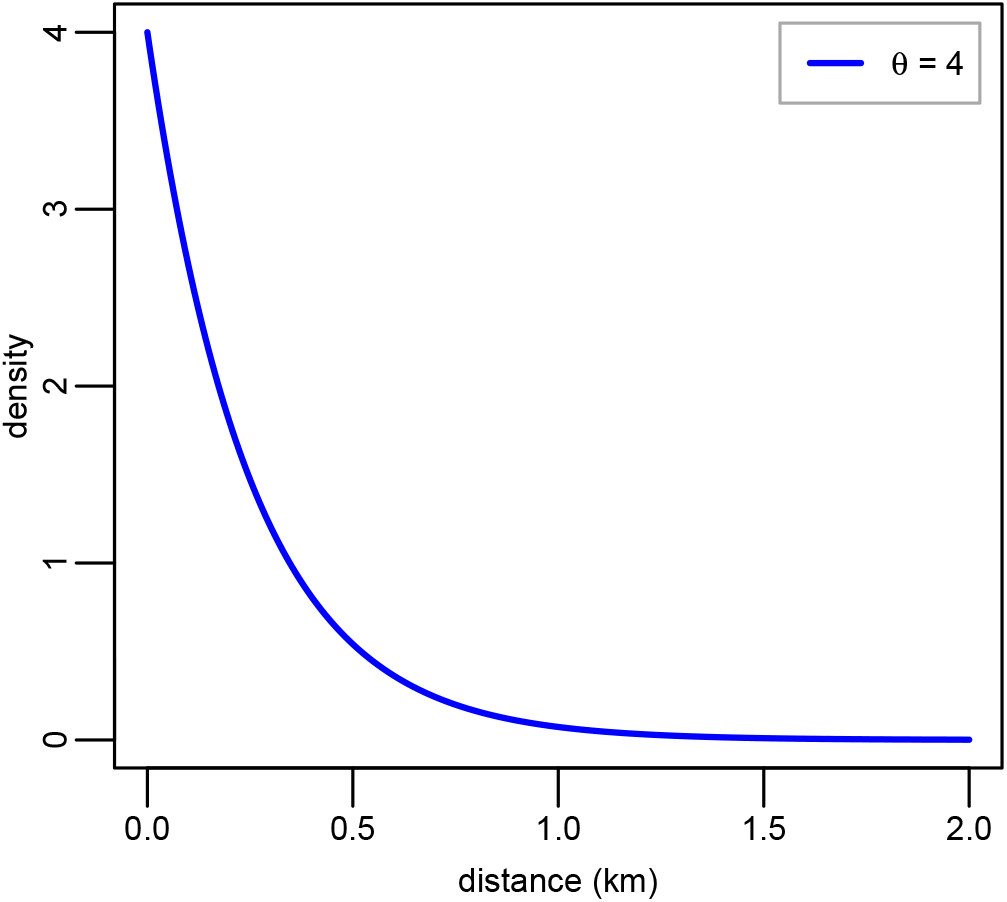
Plot of exponential distribution with *θ* = 4.

The first step to calculating **W** is to obtain the distances between each pair of hamlets (Table S2).

**Table S2:** Pairwise Euclidean distances (km) between hamlets.

|  | A | B | C | D | E | F | G | H | I | J |
| --- | --- | --- | --- | --- | --- | --- | --- | --- | --- | --- |
| A | 0.000 | 1.262 | 0.731 | 1.006 | 0.504 | 1.302 | 0.369 | 0.605 | 0.843 | 1.568 |
| B | 1.262 | 0.000 | 1.877 | 0.811 | 1.379 | 0.162 | 0.893 | 0.732 | 1.893 | 1.446 |
| C | 0.731 | 1.877 | 0.000 | 1.736 | 1.115 | 1.873 | 1.043 | 1.316 | 0.200 | 1.598 |
| D | 1.006 | 0.811 | 1.736 | 0.000 | 0.806 | 0.958 | 0.752 | 0.475 | 1.836 | 2.008 |
| E | 0.504 | 1.379 | 1.115 | 0.806 | 0.000 | 1.472 | 0.630 | 0.651 | 1.272 | 2.030 |
| F | 1.302 | 0.162 | 1.873 | 0.958 | 1.472 | 0.000 | 0.938 | 0.820 | 1.872 | 1.309 |
| G | 0.369 | 0.893 | 1.043 | 0.752 | 0.630 | 0.938 | 0.000 | 0.285 | 1.111 | 1.431 |
| H | 0.605 | 0.732 | 1.316 | 0.475 | 0.651 | 0.820 | 0.285 | 0.000 | 1.393 | 1.595 |
| I | 0.843 | 1.893 | 0.200 | 1.836 | 1.272 | 1.872 | 1.111 | 1.393 | 0.000 | 1.459 |
| J | 1.568 | 1.446 | 1.598 | 2.008 | 2.030 | 1.309 | 1.431 | 1.595 | 1.459 | 0.000 |

The next step is to apply the exponential kernel with *θ* = 4 to yield a matrix of probability densities (Table S3).

**Table S3:** Matrix of probability densities.

|  | A | B | C | D | E | F | G | H | I | J |
| --- | --- | --- | --- | --- | --- | --- | --- | --- | --- | --- |
| A | 4.000 | 0.026 | 0.215 | 0.072 | 0.534 | 0.022 | 0.913 | 0.356 | 0.138 | 0.008 |
| B | 0.026 | 4.000 | 0.002 | 0.156 | 0.016 | 2.091 | 0.112 | 0.214 | 0.002 | 0.012 |
| C | 0.215 | 0.002 | 4.000 | 0.004 | 0.046 | 0.002 | 0.062 | 0.021 | 1.794 | 0.007 |
| D | 0.072 | 0.156 | 0.004 | 4.000 | 0.159 | 0.087 | 0.197 | 0.599 | 0.003 | 0.001 |
| E | 0.534 | 0.016 | 0.046 | 0.159 | 4.000 | 0.011 | 0.322 | 0.295 | 0.025 | 0.001 |
| F | 0.022 | 2.091 | 0.002 | 0.087 | 0.011 | 4.000 | 0.094 | 0.150 | 0.002 | 0.021 |
| G | 0.913 | 0.112 | 0.062 | 0.197 | 0.322 | 0.094 | 4.000 | 1.277 | 0.047 | 0.013 |
| H | 0.356 | 0.214 | 0.021 | 0.599 | 0.295 | 0.150 | 1.277 | 4.000 | 0.015 | 0.007 |
| I | 0.138 | 0.002 | 1.794 | 0.003 | 0.025 | 0.002 | 0.047 | 0.015 | 4.000 | 0.012 |
| J | 0.008 | 0.012 | 0.007 | 0.001 | 0.001 | 0.021 | 0.013 | 0.007 | 0.012 | 4.000 |

The final step to obtain matrix W is to normalize the columns so that each column sums to 1, which conceptually means that the inhabitants of each hamlet must spend 100% of their time somewhere within the overall population (Table S4).

**Table S4:** Example matrix **W** illustrating strength of transmission from hamlet *j* (columns) to hamlet *i* (rows).

| | | Transmission from hamlet $j$ | | | | | | | | | |
| --- | --- | --- | --- | --- | --- | --- | --- | --- | --- | --- | --- |
|  |  | A | B | C | D | E | F | G | H | I | J |
| Transmission to hamlet $i$ | A | 0.637 | 0.004 | 0.035 | 0.014 | 0.099 | 0.003 | 0.130 | 0.051 | 0.023 | 0.002 |
|  | B | 0.004 | 0.603 | 0.000 | 0.030 | 0.003 | 0.323 | 0.016 | 0.031 | 0.000 | 0.003 |
|  | C | 0.034 | 0.000 | 0.650 | 0.001 | 0.009 | 0.000 | 0.009 | 0.003 | 0.297 | 0.002 |
|  | D | 0.011 | 0.024 | 0.001 | 0.758 | 0.029 | 0.013 | 0.028 | 0.086 | 0.000 | 0.000 |
|  | E | 0.085 | 0.002 | 0.008 | 0.030 | 0.739 | 0.002 | 0.046 | 0.043 | 0.004 | 0.000 |
|  | F | 0.003 | 0.315 | 0.000 | 0.016 | 0.002 | 0.617 | 0.013 | 0.022 | 0.000 | 0.005 |
|  | G | 0.145 | 0.017 | 0.010 | 0.037 | 0.059 | 0.014 | 0.568 | 0.184 | 0.008 | 0.003 |
|  | H | 0.057 | 0.032 | 0.003 | 0.113 | 0.055 | 0.023 | 0.181 | 0.577 | 0.003 | 0.002 |
|  | I | 0.022 | 0.000 | 0.292 | 0.000 | 0.005 | 0.000 | 0.007 | 0.002 | 0.663 | 0.003 |
|  | J | 0.001 | 0.002 | 0.001 | 0.000 | 0.000 | 0.003 | 0.002 | 0.001 | 0.002 | 0.980 |

Note that the extent to which a hamlet is coupled to itself depends on how peripheral it is. For example, hamlet J is distant from most other hamlets while hamlet G is more centrally located: *W*_*JJ*_ = 0.980 > *W*_*GG*_ = 0.568.

We implemented the calculation of the force of infection λ_*i*_ using matrices and vectors: In a simulation with 200 hamlets, **W** is a 200-by-200 mixing matrix and **I**_**1**_ and **I**_**2**_ are 200-by-1 column vectors. The product **W**(**I**_**1**_+**I**_**2**_) is also a 200-by-1 column vector. We then multiplied this vector entrywise by the 200-by-1 column matrix formed by *β*_*i*_/*N*_*i*_.

### Implementing the model

We implemented the model using the adaptive tau-leaping approximation to the Gillespie algorithm [3; 4; 5]. Initial conditions for the stochastic model were set at equilibria values from a deterministic version of the model [6]. All simulations were conducted in R version 3.5.1 [7].

### Simulating MDA and TTT

We assume that once a person with active or latent yaws receives curative treatment with azithromycin, the person returns to the susceptible state. We simulate a round of MDA by moving individuals in the infected or latent states back into the *S* compartment with probability equal to treatment coverage, *c*_*MDA*_. We simulate a round of TTT by first modelling screening of the population. Active cases of yaws are detected with probability *s*_*TTT*_. We assume that all detected active cases are successfully treated and therefore moved to the susceptible compartment. In hamlets in which at least 1 active case was detected, individuals with latent infections have a probability *c*_*TTT*_ of being treated and moved to the susceptible compartment. We assume that undetected active cases are untreated.

### Determining the amount of TTT required to match the impact of a round of MDA

We assess the performance of TTT by determining the number of biannual rounds of TTT required to produce the same effect as an additional round of MDA. We consider 2 different outcomes measured at 6 months following the additional round of MDA: 1) the prevalence of active infection and 2) the prevalence of latent infection. We compared TTT versus an additional round of MDA after 1, 2, or 3 initial biannual rounds of MDA. Each simulation began by generating anew the simulated geography (i.e. hamlet sizes and locations) and running the model for 100 years to establish endemic conditions in the absence of MDA or TTT. After simulating the initial rounds of MDA, we simulated an additional round of MDA in 300 parallel replicates and averaged across the replicates the prevalence of active disease and the prevalence of latent infection, both calculated 6 months following the implementation of the additional round of MDA. These prevalence estimates serve as the targets for TTT to meet or exceed. Next, we returned to the state immediately prior to implementing the additional round of MDA and began implementing biannual rounds of TTT. We calculated from 300 parallel replicates the average number of rounds of TTT and the average total number of hamlets treated in order to reach the active disease prevalence target and the latent infection prevalence target.

### Simulation experiments: Varying aspects of spatial epidemiology

We investigated how 3 different aspects of spatial epidemiology impact the relative utility of TTT versus MDA: 1) mixing between hamlets, 2) spatial heterogeneity in transmissibility, and 3) imported infections from outside the entire population. Mixing between hamlets is controlled by *θ*, which we allowed to take values of 2, 4, and 8 with larger values corresponding to less between-hamlet mixing (Figure 2A). These levels correspond to scenarios where, on average, 29%, 61%, and 86%, of contacts occur within the hamlet, respectively. (In a hypothetical scenario in which hamlets have no bearing on transmission, on average just 0.5% of contacts would occur within a hamlet.) To create spatial heterogeneity in transmission rates, hamlet-specific transmissibility coefficients *β*_*i*_ were randomly drawn from a gamma distribution with mean 5 and variance 0, 1, or 4 (Figure 2B). Finally, we varied the exogenous force of infection from outside the population by varying the rate *ξ*. We conducted simulations where *ξ* equaled zero (ie, no exogenous force of infection) and where *ξ* equaled 0.0005, which corresponds to a scenario where, in a wholly susceptible population of 20 000 people, we would expect about 10 people to become infected each year due to exposure to yaws from outside the population.

We conducted 200 simulations at each of the 18 combinations of spatial epidemiology parameters (3 levels of mixing, 3 of spatial heterogeneity of transmissibility, and 2 of exogenous force of infection). Finally, we simulated quarterly, rather than biannual, implementation of MDA and TTT for 9 combinations of spatial epidemiology parameters (3 levels of mixing and 3 levels of spatial heterogeneity of transmissibility and assuming no exogenous force of infection). For each set of 200 simulations we calculated the mean expected number of hamlets requiring treatment and the mean expected number of rounds of TTT to match the impact of the additional round of MDA (Table S5).

### Frequency with which maximum number of rounds of TTT reached

Our main goal was to ascertain the number of rounds of TTT are required to match the impact of an additional round of MDA. However, it is important to set an upper limit on the number of rounds of TTT in order to ensure that simulations do not run indefinitely. Also, from a public health programmatic perspective, extremely long TTT campaigns (e.g. lasting decades) are infeasible and therefore precisely estimating their duration is unimportant. Table S5 lists for each set of parameters the proportion of TTT campaigns that reached the 100 rounds of TTT limit. For each set of parameters and each indicator (active disease or latent infection), we simulated 60 000 TTT campaigns (300 campaigns for each of 200 baseline simulations).

For the main set of simulations with biannual interventions and no exogenous force of infection, the 100 rounds of TTT maximum was never reached. But for the simulations with an exogenous force of infection, the maximum number of rounds of TTT was reached in some replicates when the target indicator was prevalence of latent infection and there were 2 or 3 initial rounds of MDA. The corresponding mean number of rounds of TTT and mean hamlets treated therefore are underestimates. For this reason, we do not plot these means in the main text. Instead, we calculated the probability of matching the impact of the additional round of MDA as a function of number of rounds of TTT (Figure 5). For simulations with quarterly interventions and when the target indicator was prevalence of latent infection, the 100 rounds of TTT maximum was reached so rarely (a total of 13 times out of 1.62 million simulated TTT campaigns) it would not have a meaningful impact on our results.

**Table S5:** Summary of results for each set of parameters, by target indicator (prevalence of active disease or prevalence of latent infection). Entries marked with a strikethrough indicates that the 100 rounds of TTT maximum was reached at least once for that set of parameters.

| $f$ | $\xi$ | $\beta_{var}$ | $\theta$ | initial<br>MDA<br>rounds | Indicator: prevalence of<br>active disease | | | Indicator: prevalence of<br>latent infection | | |
| --- | --- | --- | --- | --- | --- | --- | --- | --- | --- | --- |
| | | | | | mean<br>TTT<br>rounds | mean<br>hamlets<br>treated | $\geq 100$<br>rounds<br>TTT | mean<br>TTT<br>rounds | mean<br>hamlets<br>treated | $\geq 100$<br>rounds<br>TTT |
| 0.5 | 0 | 0 | 2 | 1 | 3.44 | 111.03 | 0 | 7.85 | 141.44 | 0 |
| 0.5 | 0 | 0 | 2 | 2 | 2.70 | 17.17 | 0 | 10.78 | 25.72 | 0 |
| 0.5 | 0 | 0 | 2 | 3 | 1.31 | 2.46 | 0 | 9.00 | 3.75 | 0 |
| 0.5 | 0 | 0 | 4 | 1 | 3.57 | 84.05 | 0 | 8.00 | 106.85 | 0 |
| 0.5 | 0 | 0 | 4 | 2 | 2.58 | 11.93 | 0 | 9.90 | 17.58 | 0 |
| 0.5 | 0 | 0 | 4 | 3 | 1.27 | 1.58 | 0 | 8.31 | 2.40 | 0 |
| 0.5 | 0 | 0 | 8 | 1 | 3.41 | 39.56 | 0 | 8.14 | 51.63 | 0 |
| 0.5 | 0 | 0 | 8 | 2 | 2.06 | 4.72 | 0 | 9.76 | 7.38 | 0 |
| 0.5 | 0 | 0 | 8 | 3 | 1.14 | 0.48 | 0 | 6.29 | 0.80 | 0 |
| 0.5 | 0 | 1 | 2 | 1 | 3.28 | 108.92 | 0 | 7.65 | 139.88 | 0 |
| 0.5 | 0 | 1 | 2 | 2 | 2.72 | 17.97 | 0 | 10.68 | 26.93 | 0 |
| 0.5 | 0 | 1 | 2 | 3 | 1.32 | 2.18 | 0 | 9.18 | 3.65 | 0 |
| 0.5 | 0 | 1 | 4 | 1 | 3.12 | 81.98 | 0 | 7.44 | 107.33 | 0 |
| 0.5 | 0 | 1 | 4 | 2 | 2.38 | 12.89 | 0 | 9.31 | 19.25 | 0 |
| 0.5 | 0 | 1 | 4 | 3 | 1.30 | 1.91 | 0 | 7.87 | 2.84 | 0 |
| 0.5 | 0 | 1 | 8 | 1 | 2.72 | 46.41 | 0 | 6.87 | 63.32 | 0 |
| 0.5 | 0 | 1 | 8 | 2 | 2.06 | 7.33 | 0 | 8.19 | 11.03 | 0 |
| 0.5 | 0 | 1 | 8 | 3 | 1.24 | 0.97 | 0 | 7.07 | 1.52 | 0 |
| 0.5 | 0 | 4 | 2 | 1 | 2.86 | 103.05 | 0 | 7.17 | 136.11 | 0 |
| 0.5 | 0 | 4 | 2 | 2 | 2.50 | 19.37 | 0 | 10.10 | 29.31 | 0 |
| 0.5 | 0 | 4 | 2 | 3 | 1.36 | 2.75 | 0 | 9.42 | 4.49 | 0 |
| 0.5 | 0 | 4 | 4 | 1 | 2.38 | 81.22 | 0 | 6.01 | 112.02 | 0 |
| 0.5 | 0 | 4 | 4 | 2 | 2.07 | 17.87 | 0 | 7.17 | 26.38 | 0 |
| 0.5 | 0 | 4 | 4 | 3 | 1.49 | 4.30 | 0 | 7.02 | 5.90 | 0 |
| 0.5 | 0 | 4 | 8 | 1 | 2.24 | 54.95 | 0 | 4.95 | 76.11 | 0 |
| 0.5 | 0 | 4 | 8 | 2 | 2.00 | 14.05 | 0 | 4.53 | 19.16 | 0 |
| 0.5 | 0 | 4 | 8 | 3 | 1.70 | 4.80 | 0 | 4.29 | 5.63 | 0 |
| 0.5 | 0.0005 | 0 | 2 | 1 | 3.26 | 122.79 | 0 | 8.12 | 176.67 | 0 |
| 0.5 | 0.0005 | 0 | 2 | 2 | 2.62 | 31.24 | 0 | ~~42.04~~ | ~~254.61~~ | 6680 |
| 0.5 | 0.0005 | 0 | 2 | 3 | 2.04 | 13.55 | 0 | ~~80.00~~ | ~~436.24~~ | 41812 |
| 0.5 | 0.0005 | 0 | 4 | 1 | 3.46 | 95.34 | 0 | 8.30 | 137.46 | 0 |

Table S5(Continued)
|  |  |  |  |  |  |  |  |  |  |  |
| --- | --- | --- | --- | --- | --- | --- | --- | --- | --- | --- |
| 0.5 | 0.0005 | 0 | 4 | 2 | 2.46 | 21.42 | 0 | 31.04 | 149.61 | 2384 |
| 0.5 | 0.0005 | 0 | 4 | 3 | 1.96 | 10.15 | 0 | 59.93 | 257.54 | 25489 |
| 0.5 | 0.0005 | 0 | 8 | 1 | 3.40 | 55.95 | 0 | 8.96 | 88.74 | 0 |
| 0.5 | 0.0005 | 0 | 8 | 2 | 2.25 | 13.11 | 0 | 30.53 | 112.41 | 2682 |
| 0.5 | 0.0005 | 0 | 8 | 3 | 1.89 | 7.32 | 0 | 42.84 | 146.37 | 14219 |
| 0.5 | 0.0005 | 1 | 2 | 1 | 3.09 | 120.83 | 0 | 7.87 | 175.23 | 0 |
| 0.5 | 0.0005 | 1 | 2 | 2 | 2.66 | 31.10 | 0 | 42.54 | 258.09 | 6955 |
| 0.5 | 0.0005 | 1 | 2 | 3 | 2.04 | 14.09 | 0 | 79.61 | 438.25 | 41405 |
| 0.5 | 0.0005 | 1 | 4 | 1 | 3.04 | 92.57 | 0 | 7.69 | 135.93 | 0 |
| 0.5 | 0.0005 | 1 | 4 | 2 | 2.40 | 22.89 | 0 | 25.13 | 127.50 | 1149 |
| 0.5 | 0.0005 | 1 | 4 | 3 | 1.94 | 11.14 | 0 | 57.34 | 250.25 | 22562 |
| 0.5 | 0.0005 | 1 | 8 | 1 | 2.77 | 57.97 | 0 | 7.38 | 90.86 | 0 |
| 0.5 | 0.0005 | 1 | 8 | 2 | 2.18 | 15.26 | 0 | 19.04 | 77.12 | 513 |
| 0.5 | 0.0005 | 1 | 8 | 3 | 1.89 | 8.12 | 0 | 39.06 | 135.57 | 11282 |
| 0.5 | 0.0005 | 4 | 2 | 1 | 2.83 | 112.32 | 0 | 7.49 | 167.33 | 0 |
| 0.5 | 0.0005 | 4 | 2 | 2 | 2.44 | 31.69 | 0 | 37.07 | 233.22 | 5347 |
| 0.5 | 0.0005 | 4 | 2 | 3 | 2.01 | 14.70 | 0 | 75.72 | 426.25 | 38186 |
| 0.5 | 0.0005 | 4 | 4 | 1 | 2.37 | 88.82 | 0 | 6.20 | 134.08 | 0 |
| 0.5 | 0.0005 | 4 | 4 | 2 | 2.14 | 26.50 | 0 | 14.19 | 89.12 | 198 |
| 0.5 | 0.0005 | 4 | 4 | 3 | 1.92 | 12.97 | 0 | 38.73 | 180.57 | 12829 |
| 0.5 | 0.0005 | 4 | 8 | 1 | 2.23 | 61.72 | 0 | 5.15 | 92.65 | 0 |
| 0.5 | 0.0005 | 4 | 8 | 2 | 2.04 | 21.12 | 0 | 6.59 | 42.71 | 6 |
| 0.5 | 0.0005 | 4 | 8 | 3 | 1.94 | 11.52 | 0 | 15.37 | 60.42 | 3059 |
| 0.25 | 0 | 0 | 2 | 1 | 3.00 | 86.88 | 0 | 13.52 | 131.63 | 0 |
| 0.25 | 0 | 0 | 2 | 2 | 2.47 | 11.79 | 0 | 18.76 | 21.06 | 0 |
| 0.25 | 0 | 0 | 2 | 3 | 1.19 | 1.16 | 0 | 15.59 | 2.13 | 1 |
| 0.25 | 0 | 0 | 4 | 1 | 3.20 | 69.18 | 0 | 14.16 | 107.13 | 0 |
| 0.25 | 0 | 0 | 4 | 2 | 2.36 | 8.69 | 0 | 18.47 | 15.75 | 0 |
| 0.25 | 0 | 0 | 4 | 3 | 1.18 | 0.88 | 0 | 14.25 | 1.60 | 5 |
| 0.25 | 0 | 0 | 8 | 1 | 3.11 | 34.51 | 0 | 14.63 | 55.41 | 0 |
| 0.25 | 0 | 0 | 8 | 2 | 1.88 | 4.11 | 0 | 18.02 | 7.67 | 0 |
| 0.25 | 0 | 0 | 8 | 3 | 1.09 | 0.37 | 0 | 10.37 | 0.66 | 0 |
| 0.25 | 0 | 1 | 2 | 1 | 2.94 | 84.91 | 0 | 13.38 | 128.87 | 0 |
| 0.25 | 0 | 1 | 2 | 2 | 2.39 | 11.73 | 0 | 18.66 | 21.02 | 0 |
| 0.25 | 0 | 1 | 2 | 3 | 1.20 | 1.24 | 0 | 15.43 | 2.25 | 2 |
| 0.25 | 0 | 1 | 4 | 1 | 2.91 | 68.06 | 0 | 13.52 | 106.23 | 0 |
| 0.25 | 0 | 1 | 4 | 2 | 2.26 | 9.62 | 0 | 17.84 | 16.99 | 0 |
| 0.25 | 0 | 1 | 4 | 3 | 1.20 | 1.22 | 0 | 14.39 | 1.96 | 1 |
| 0.25 | 0 | 1 | 8 | 1 | 2.62 | 41.87 | 0 | 12.95 | 67.57 | 0 |
| 0.25 | 0 | 1 | 8 | 2 | 2.00 | 6.26 | 0 | 17.11 | 11.23 | 0 |
| 0.25 | 0 | 1 | 8 | 3 | 1.16 | 0.82 | 0 | 12.62 | 1.34 | 0 |
| 0.25 | 0 | 4 | 2 | 1 | 2.67 | 80.25 | 0 | 12.90 | 123.18 | 0 |
| 0.25 | 0 | 4 | 2 | 2 | 2.22 | 12.49 | 0 | 18.31 | 21.77 | 0 |
| 0.25 | 0 | 4 | 2 | 3 | 1.22 | 1.54 | 0 | 15.70 | 2.61 | 2 |
| 0.25 | 0 | 4 | 4 | 1 | 2.34 | 65.86 | 0 | 12.19 | 105.25 | 0 |
| 0.25 | 0 | 4 | 4 | 2 | 2.07 | 10.82 | 0 | 16.79 | 19.21 | 0 |
| 0.25 | 0 | 4 | 4 | 3 | 1.23 | 1.43 | 0 | 14.23 | 2.36 | 2 |
| 0.25 | 0 | 4 | 8 | 1 | 2.12 | 45.52 | 0 | 10.96 | 74.48 | 0 |
| 0.25 | 0 | 4 | 8 | 2 | 1.86 | 8.45 | 0 | 14.77 | 15.04 | 0 |
| 0.25 | 0 | 4 | 8 | 3 | 1.29 | 1.58 | 0 | 12.92 | 2.35 | 0 |

### Comparing the cost of TTT versus the cost of MDA

We consulted published literature on yaws interventions as well as on other mass drug administration campaigns to develop illustrative costing scenarios [8; 9]. The cost of implementing MDA in various settings is relatively well documented, but TTT-related costs are more speculative. In the low-operational-cost scenario, we assume that the population is relatively easy to access and that the cost of the drug itself is a comparatively large expense, whereas in the high-operational-cost scenario, we assume that accessing the population is challenging and therefore logistical costs are high (Table S6).

**Table S6:** Indicative costs (USD) for implementing MDA and TTT in a low-operational-cost scenario and in a high-operational-cost scenario.

| Parameter | Description of expense | Low-operational-cost scenario (USD) | High-operational-cost scenario (USD) |
| --- | --- | --- | --- |
| $x_{drug}$ | drug (per person) | 0.80 | 0.80 |
| $x_{MDA}$ | MDA excluding drug (per person per round) | 1.50 | 6.00 |
| $x_{TTT_{screen}}$ | TTT/screening excluding drug (per person per round)<sup>1</sup> | 0.70 | 2.80 |
| $x_{TTT_{treat}}$ | TTT/drug admin excluding drug (per person per round)<sup>2</sup> | 0.80 | 3.20 |

<sup>1</sup>This is the cost per person in the entire population, not just among those who are identified as cases and contacts.
<sup>2</sup>This is the cost per person among those who are identified as cases and contacts during TTT.

We calculated in terms of *r*, the number of rounds of TTT, and *h*, the total hamlets treated, the threshold where the cost of a TTT campaign matches the cost of a round of MDA. First, we calculated an expression for the cost of a round of MDA:

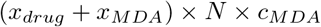

And an expression for the cost of a TTT campaign:

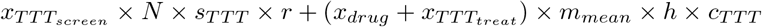

This expression makes the following simplifying assumptions: It uses *m*_*mean*_ rather than the specific values for *m* of the hamlets that received treatment and it assumes that *c*_*TTT*_ equals *s*_*TTT*_.

We find the threshold at which TTT and MDA are equivalent by setting these expressions equal:

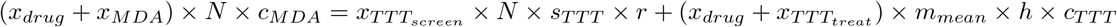

In our simulations *c*_*MDA*_ = *c*_*TTT*_ = *s*_*TTT*_, so these parameters can be factored out:

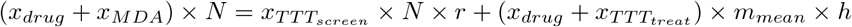

We can substitute in *N/n* for *m*_*mean*_:

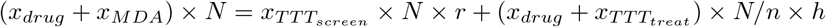

And then factor out *N*:

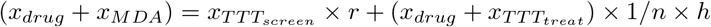

Finally, we solve for h in terms of r:

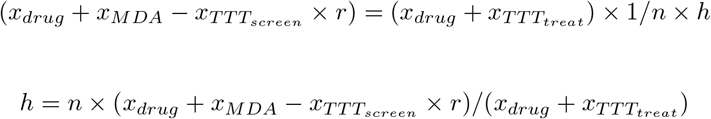

For the low operation-cost scenario we obtain the following equation:

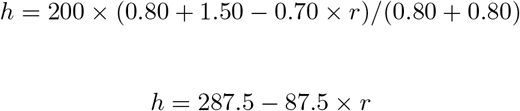

Under the low-operational-cost scenario, 1 round of MDA is economically equivalent to 2 rounds of TTT in which the total number of people treated is 56.25% of the number that would be treated in a single round of MDA and is equivalent to 3 rounds of TTT in which the total number of people treated is 12.5% of the number that would be treated in a single round of MDA. If it took on average more than 3.29 rounds of TTT to equal the impact of 1 additional round of MDA, TTT would not be economically preferable, no matter how few people had to be treated.

For the high operation-cost scenario we obtain the following equation:

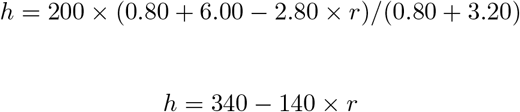

Under the high-operational-cost scenario, 1 round of MDA is economically equivalent to 2 rounds of TTT in which the total number of people treated is 30% of the number that would be treated in a single round of MDA. If it took on average more than 2.43 rounds of TTT to equal the impact of 1 additional round of MDA, TTT would not be economically preferable, no matter how few people had to be treated.

### Impact of changing spatial epidemiology parameters on performance of TTT versus MDA

For ease of interpretation, Figure 3 only shows for an intermediate level of mixing (*θ* = 4) and an intermediate level of heterogeneity in transmissibility (*β*_*var*_ = 1) the consequence of using active disease versus latent infection as the indicator of intervention impact. Figure S4 shows results for both indicators and all 9 combinations of *θ* and *β*_*var*_.

**Figure S4:**
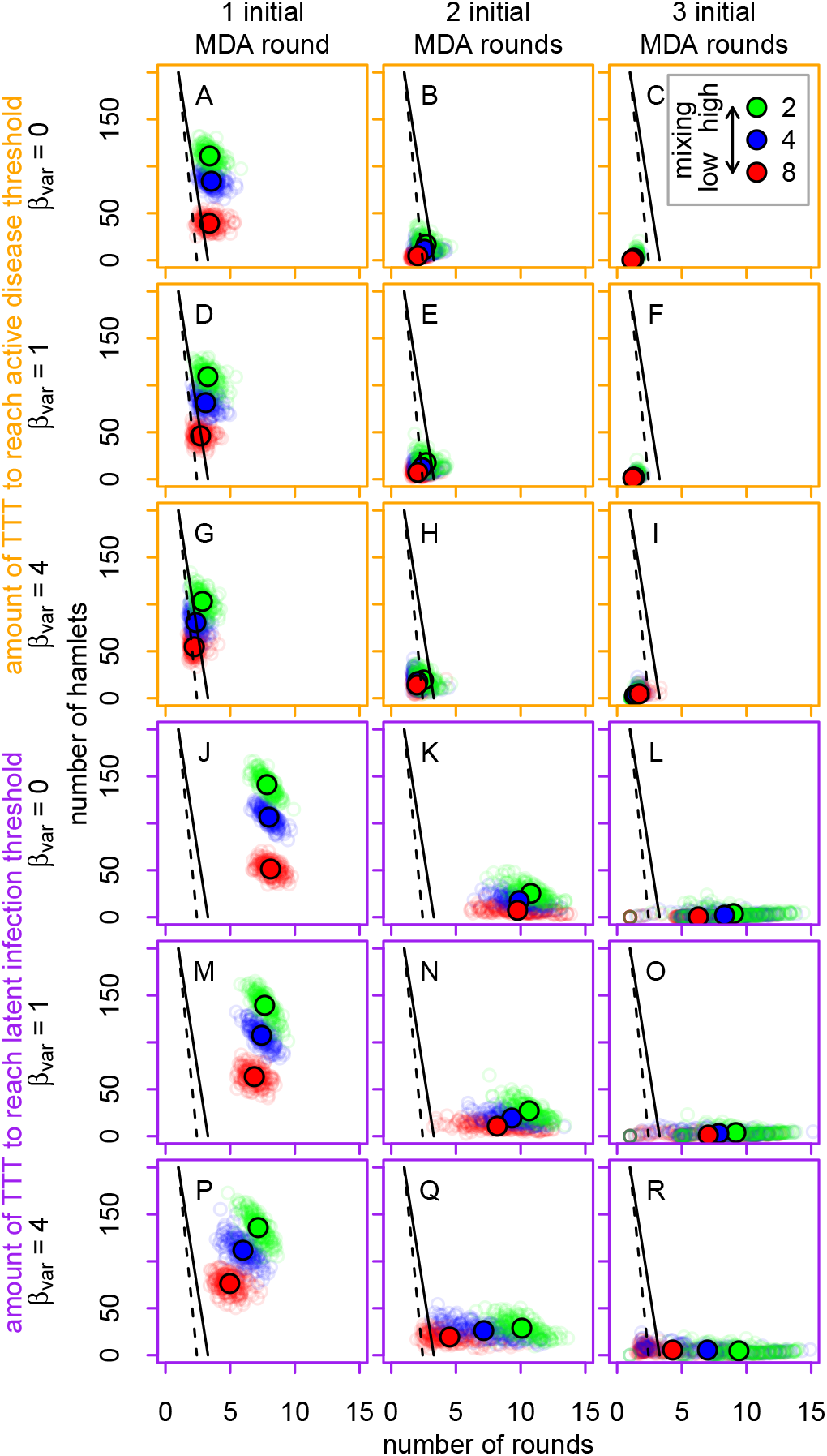
Number of rounds of TTT and total number of hamlets that must be treated in order to match the impact of a single round of MDA. The indicator of intervention impact is prevalence of active disease in the panels with orange borders (A-I) and is prevalence of latent infection in the panels with purple borders (J-R). The colors of the points correspond to mixing between hamlets, the rows to spatial heterogeneity in transmissibility, and the columns to the initial number of rounds of MDA. The large solid colored circles mark the mean result for each set of parameters, while the small circles correspond to means calculated from different baseline states. The solid and dashed black lines correspond to the thresholds where TTT and MDA cost the same and represent low- and high-operational-cost scenarios, respectively.

### Stochastic variability in eradication campaigns and the precision of estimates

The main source of variability in Figure S4 is variability in baseline states (i.e. stochastic variability in the endemic equilibrium, in the impact of the initial round(s) of MDA, and in transmission between rounds of MDA and for 6 months immediately after the last initial round of MDA), rather than uncertainty in the average amount of TTT required to match the average impact of an additional round of MDA (Figures S5 and S6). The actual (as opposed to mean) amount of TTT required to match the average impact of an additional round of MDA varies widely between replicates, especially when prevalence of yaws is very low (Figures S7 and S8).

**Figure S5:**
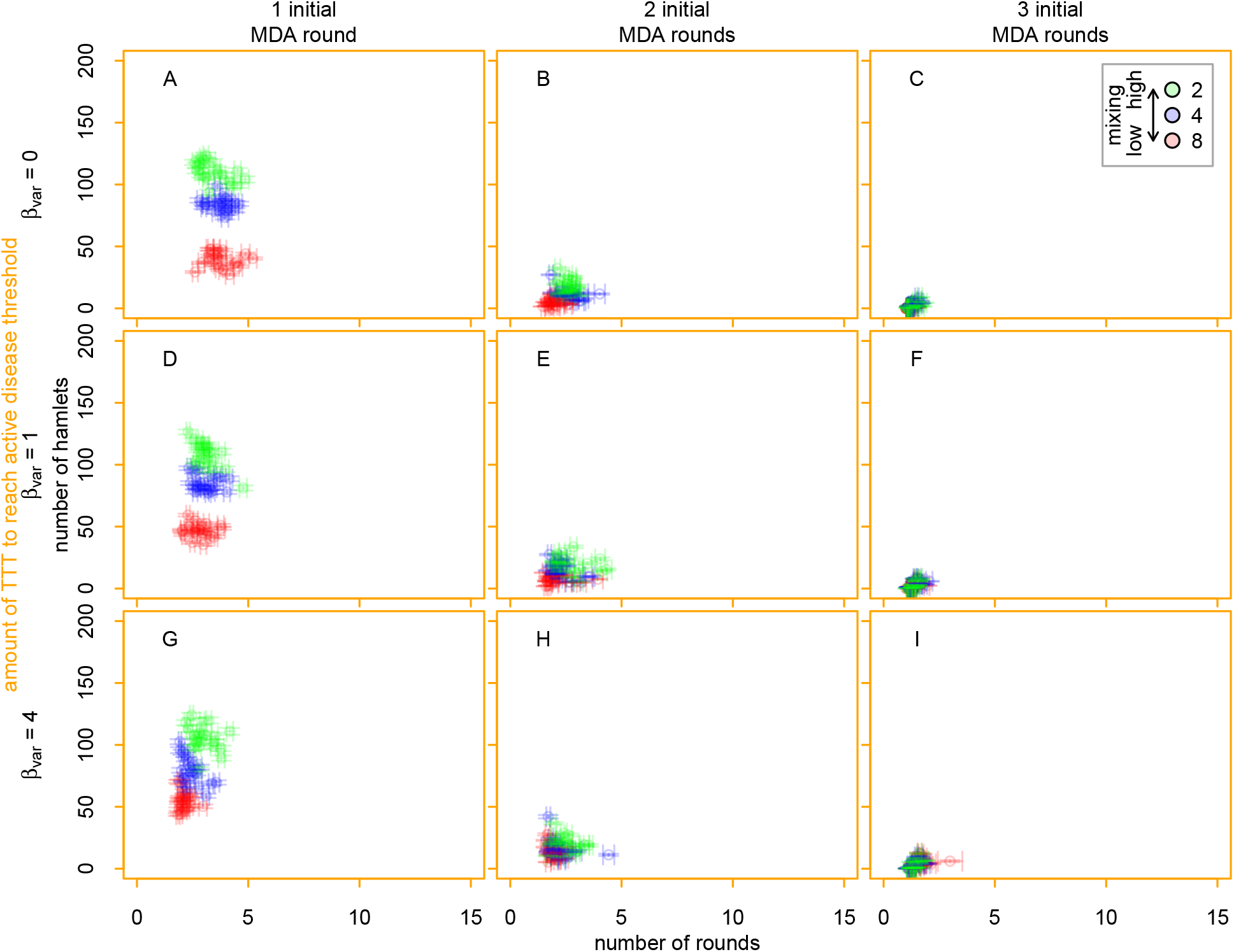
Precision of estimates for each baseline state of the number of rounds of TTT and the total number of hamlets that must be treated in order to match the impact of a single round of MDA, as defined by the magnitude of the reduction in the prevalence of active disease. The colors of the points correspond to mixing between hamlets, the rows to spatial heterogeneity in transmissibility, and the columns to the initial number of rounds of MDA. The error bars represent 95% Wald confidence intervals about the mean number of rounds of TTT (horizontal bars) or mean total amount of treatment (vertical bars) required to match the mean impact of a single round of MDA, for a given baseline state. For ease of display, only 20 sets of baseline states are shown for each combination of parameters.

**Figure S6:**
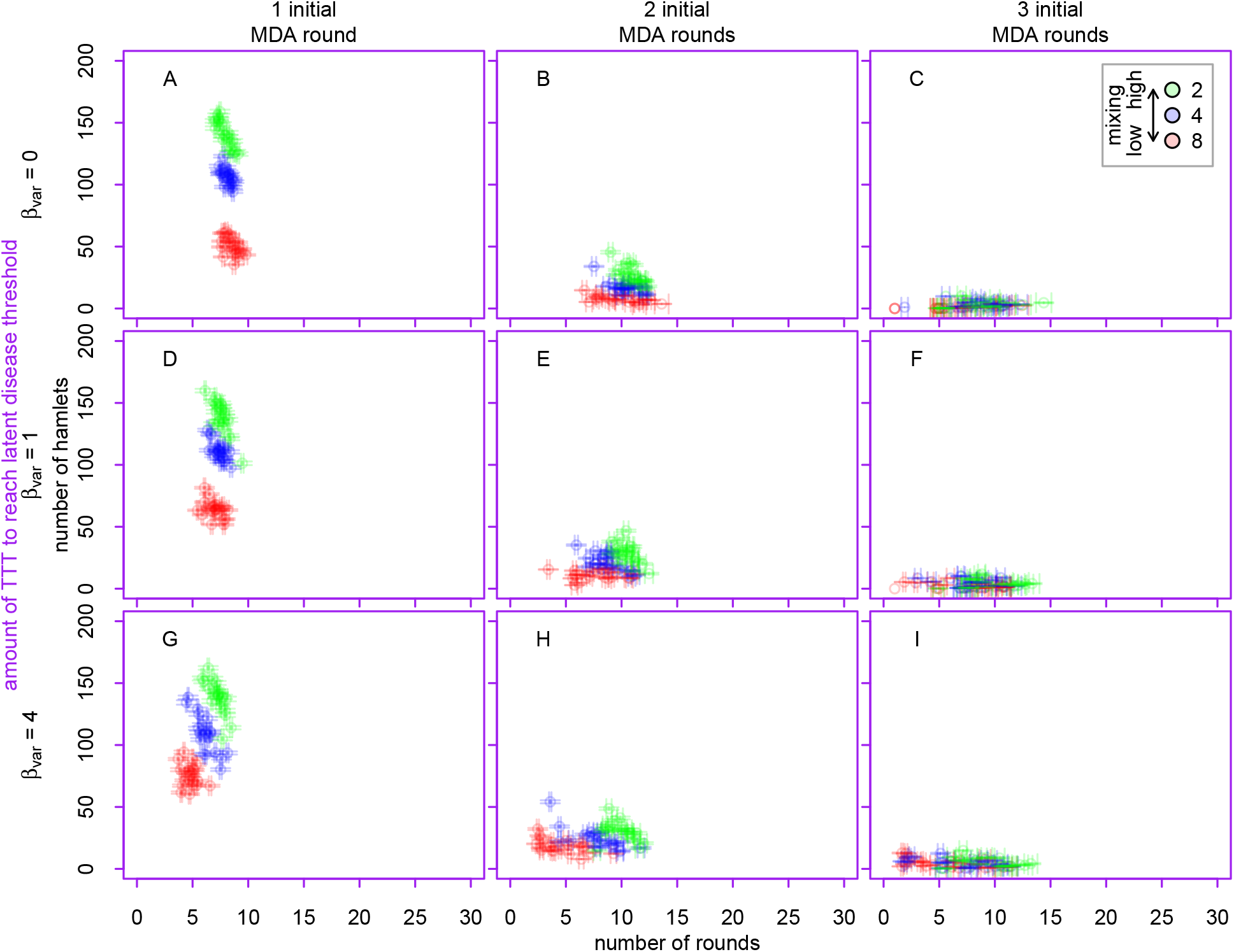
Precision of estimates for each baseline state of the number of rounds of TTT and the total number of hamlets that must be treated in order to match the impact of a single round of MDA, as defined by the magnitude of the reduction in the prevalence of latent infection. The colors of the points correspond to mixing between hamlets, the rows to spatial heterogeneity in transmissibility, and the columns to the initial number of rounds of MDA. The error bars represent 95% Wald confidence intervals about the mean number of rounds of TTT (horizontal bars) or mean total amount of treatment (vertical bars) required to match the mean impact of a single round of MDA, for a given baseline state. For ease of display, only 20 sets of baseline states are shown for each combination of parameters.

**Figure S7:**
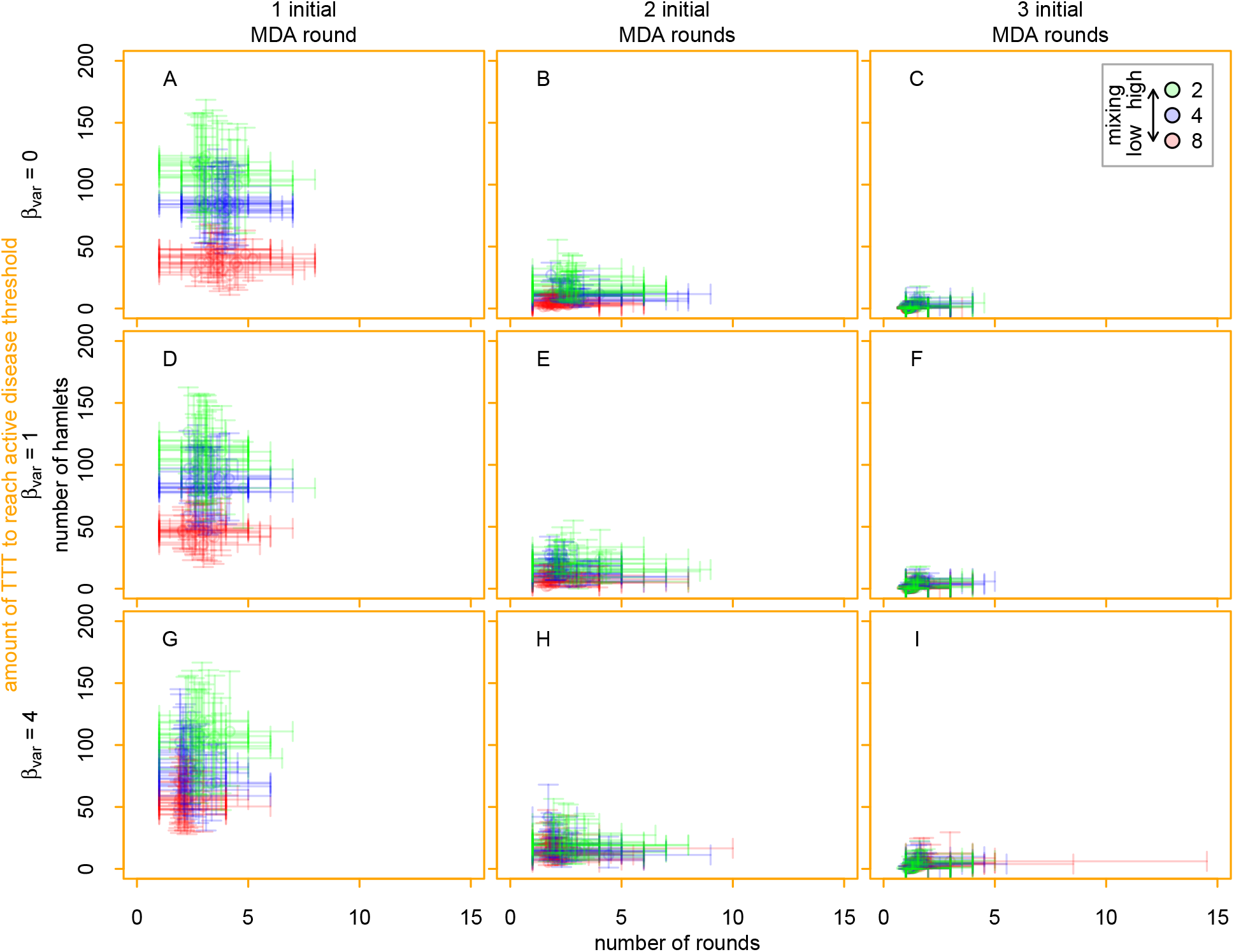
Variability for each baseline state in the number of rounds of TTT and the total number of hamlets that must be treated across those rounds in order to match the impact of a single round of MDA, as defined by the magnitude of the reduction in the prevalence of active disease. The colors of the points correspond to mixing between hamlets, the rows to spatial heterogeneity in transmissibility, and the columns to the initial number of rounds of MDA. The error bars represent the underlying variability in the 300 replicates simulated from each baseline state. The bars mark the 2.5th to 97.5th percentiles of the number of rounds of TTT (horizontal bars) or total amount of treatment (vertical bars) required to match the mean impact of a single round of MDA. For ease of display, only 20 sets of baseline states are shown for each combination of parameters.

**Figure S8:**
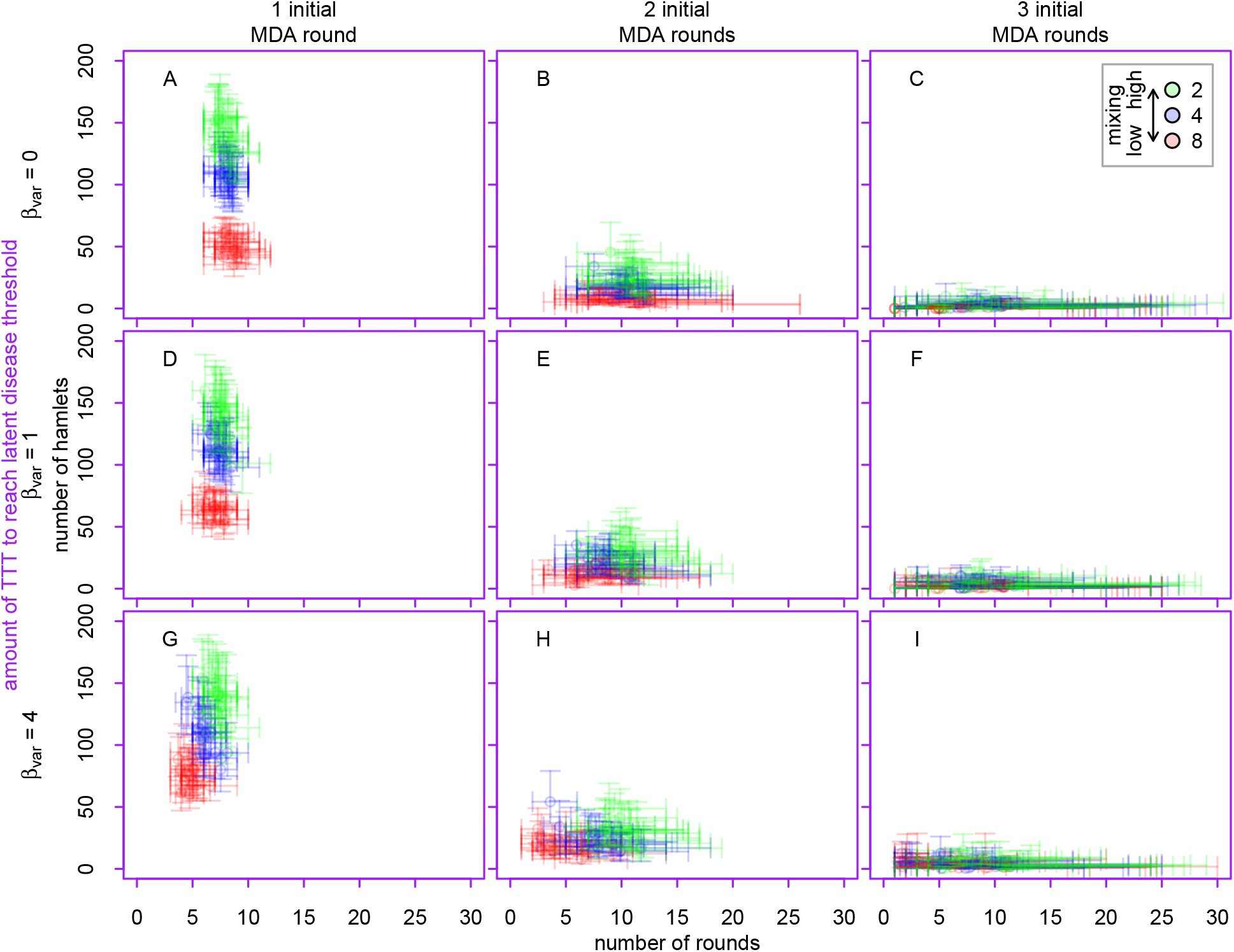
Variability for each baseline state in the number of rounds of TTT and the total number of hamlets that must be treated across those rounds in order to match the impact of a single round of MDA, as defined by the magnitude of the reduction in the prevalence of latent infection. The colors of the points correspond to mixing between hamlets, the rows to spatial heterogeneity in transmissibility, and the columns to the initial number of rounds of MDA. The error bars represent the underlying variability in the 300 replicates simulated from each baseline state. The bars mark the 2.5th to 97.5th percentiles of the number of rounds of TTT (horizontal bars) or total amount of treatment (vertical bars) required to match the mean impact of a single round of MDA. For ease of display, only 20 sets of baseline states are shown for each combination of parameters.

### Performance of TTT when yaws is imported from outside the study area

**Figure S9:**
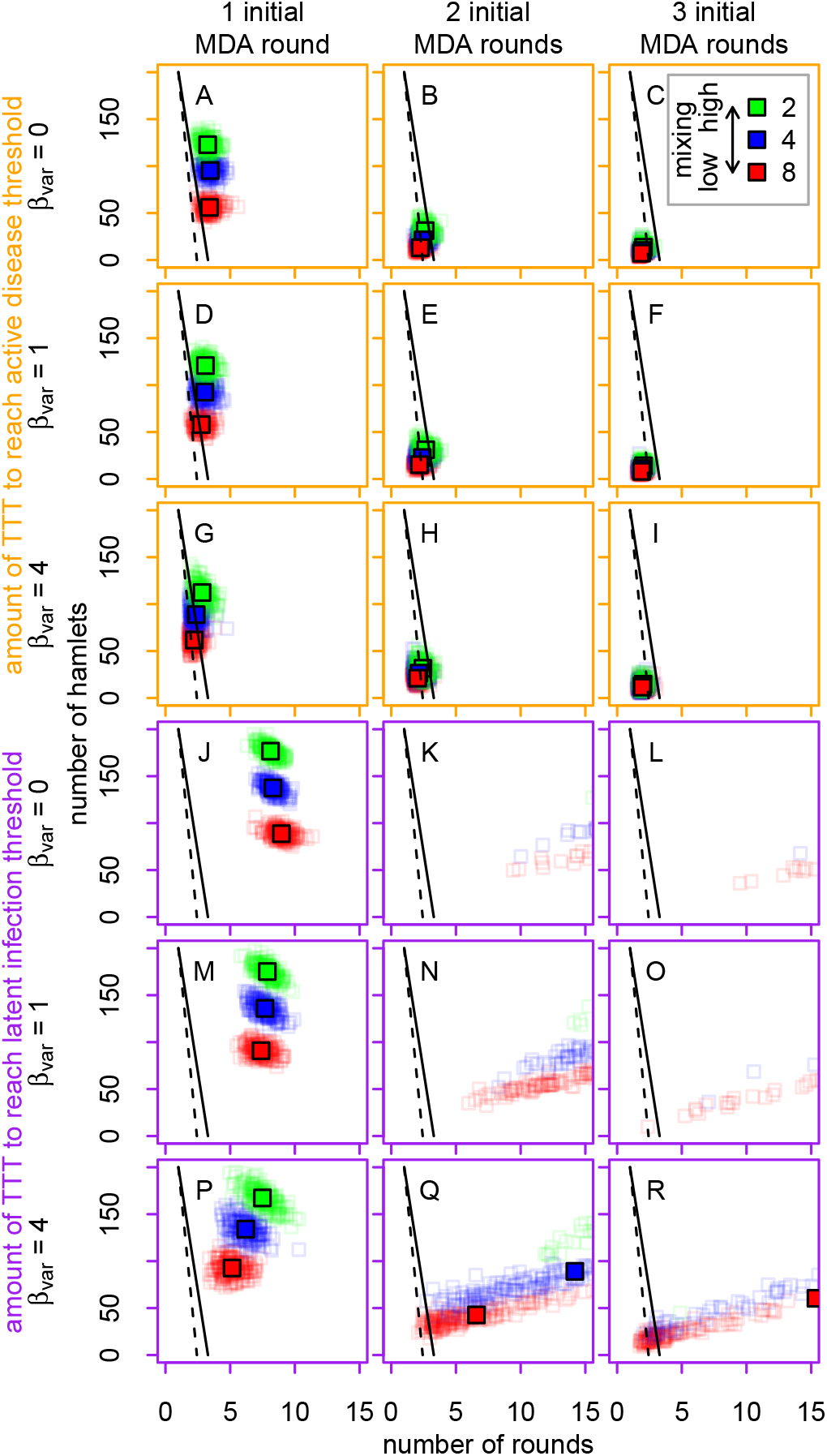
Performance of TTT versus MDA in a setting with an exogenous force of infection at a rate 0.0005 years^−1^. This figure plots the number of rounds of TTT and the total number of hamlets that must be treated across those rounds in order to match the impact of a single round of MDA, as defined by the magnitude of the reduction in the prevalence of active disease (orange panels, A-I) or latent infection (purple panels, J-R). The colors of the points correspond to mixing between hamlets, the rows to spatial heterogeneity in transmissibility, and the columns to the initial number of rounds of MDA. The large solid colored squares mark the mean result for each set of parameters, while the small squares reflect stochastic variability in initial conditions and in the impact of the initial rounds of MDA. Panels K, N, L, O, Q, and R are affected by the 100 rounds of TTT maximum (Table S5). The solid and dashed black lines correspond to the thresholds where TTT and MDA cost the same and represent low- and high-operational-cost scenarios, respectively.

### Changing the frequency of MDA and TTT to be quarterly

**Figure S10:**
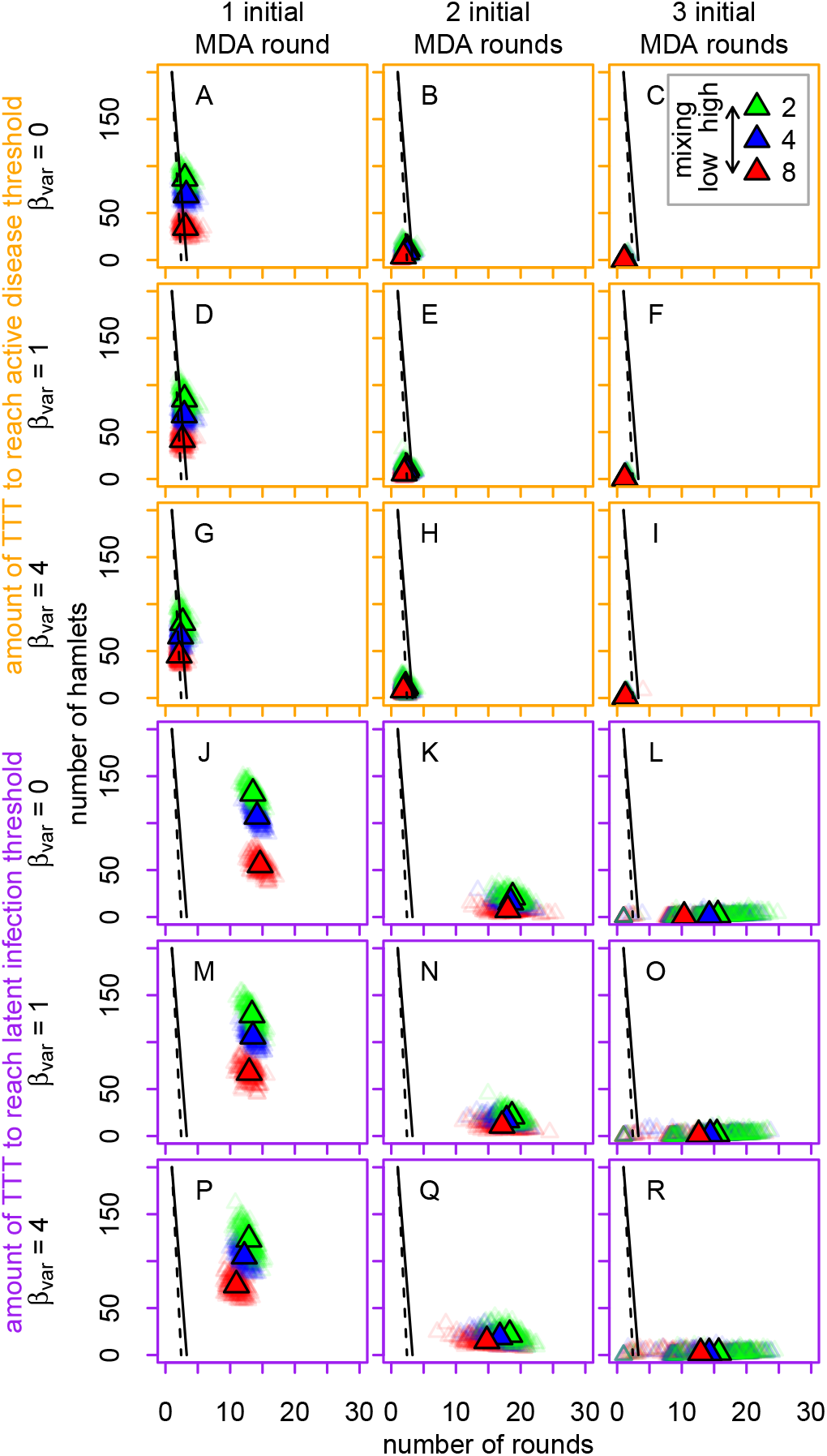
Performance of TTT versus MDA when MDA and TTT are implemented quarterly. This figure plots the number of rounds of TTT and the total number of hamlets that must be treated across those rounds in order to match the impact of a single round of MDA, as defined by the magnitude of the reduction in the prevalence of active disease (orange panels, A-I) or latent infection (purple panels, J-R). The colors of the points correspond to mixing between hamlets, the rows to spatial heterogeneity in transmissibility, and the columns to the initial number of rounds of MDA. The large solid colored triangles mark the mean result for each set of parameters, while the small triangles reflect stochastic variability in initial conditions and in the impact of the initial rounds of MDA. The solid and dashed black lines correspond to the thresholds where TTT and MDA cost the same and represent low- and high-operational-cost scenarios, respectively. Note that the x-axis ranges from 0 to 30.

